# Asymmetric partition of the *O*-GlcNAcome in mitosis ensures binary cell fate decision

**DOI:** 10.64898/2026.01.18.700201

**Authors:** Fang Chen, Haibin Yu, Wanxin Zeng, Xiaoling Wei, Song Mao, Lu Lv, Hongtao Qin, Ke Liu, Hongda Huang, Zhuohua Zhang, Xuebiao Yao, Kai Yuan

**Affiliations:** Hunan Key Laboratory of Molecular Precision Medicine, Department of Oncology, Xiangya Hospital & Center for Medical Genetics, School of Life Sciences, Central South University, Changsha, China; National Clinical Research Center for Geriatric Disorders, Xiangya Hospital, Central South University, Changsha, China; State Key Laboratory of Chemo/Biosensing and Chemometrics, College of Biology, Hunan University, Changsha, China; Hubei Key Laboratory of Genetic Regulation and Integrative Biology, School of Life Sciences, Central China Normal University, Wuhan, China; Department of Chemical Biology, School of Life Sciences, Southern University of Science and Technology, Shenzhen, China; MOE Key Laboratory for Membraneless Organelles and Cellular Dynamics, Hefei National Center for Cross-disciplinary Sciences, University of Science and Technology of China, Hefei, China; The Biobank of Xiangya Hospital, Central South University, Changsha, China; Furong Laboratory, Changsha, China

**Author notes:** Correspondence (K.Y.). These authors contributed equally (Fang Chen, Haibin Yu).

**Keywords:** *O*-GlcNAcylation, 14-3-3, Nuclear pore complex, Cell fate determination, Brain development

## Abstract

Many cellular components undergo biased segregation during stem cell division, but whether such order extends to the proteome has remained unknown. Using our real-time *O*-GlycoTracer, we find that *O*-GlcNAcylated proteins (the *O*-GlcNAcome) segregate asymmetrically during *Drosophila* neuroblast mitosis, predisposing the two daughters to distinct fates. The daughter that preserves stem cell identity inherits most of the *O*-GlcNAcome. This asymmetric partition requires putative *O*-GlcNAc readers, notably 14-3-3 proteins. We further identify the nuclear pore complex (NPC) as a major *O*-GlcNAc substrate in neuroblasts; Nup153, located in the nuclear basket, shows *O*-GlcNAc-dependent biased inheritance. Perturbing multiple steps in this segregation pathway disrupts neuroblast differentiation, reduces brain size, and causes adult learning deficits. These findings reveal a coordinated *O*-GlcNAc-driven mechanism for proteome-level asymmetric inheritance during neural stem cell division, with implications for neurodevelopmental disorders linked to OGT or 14-3-3 mutations.

## Introduction

The asymmetry in cell division cycles underpins the unique self-renewal capacity of stem cells. Inherent cellular polarity, together with extrinsic cues from the stem cell niche, drives biased allocation of many cellular components between the two daughter cells during mitosis, so that one progeny perpetuates the attributes of stemness while the other becomes committed to generate diverse cell types of different functions. During the asymmetric cell divisions, fate-determining transcription regulators^1–4^, signaling molecules^5,6^, subcellular organelles^7–13^, and even certain histone modifications^14,15^ have been reported to be unequally partitioned, triggering the bifurcation of cell fates. In addition, asymmetric cell divisions have been hypothesized to promote stem cell rejuvenation by retaining a fraction of the original DNA strands^16,17^ and degradative machinery^11^ in the self-renewing daughter cell, meanwhile segregating away damaged proteins^18,19^ and organelles such as mitochondria^20^ into the differentiating counterpart.

Thousands of cellular proteins are subject to *O*-GlcNAcylation^21^, a reversible post-translational modification consisting of a single N-acetylglucosamine moiety attached to serine or threonine residues, which is catalyzed by two mutually antagonistic enzymes, *O*-GlcNAc transferase (OGT) and *O*-GlcNAcase (OGA)^22^. Coupled to cellular metabolic plasticity, *O*-GlcNAcylation is critical for the self-renewal of stem cells, and reduction of global *O*-GlcNAcylation level promotes differentiation^23,24^. Consistently, *OGT* mutations or *O*-GlcNAcylation deficiencies result in developmental defects and early embryonic lethality in multiple model organisms^25–27^. In humans, genetic studies have established a strong link between *OGT* mutations and neurodevelopmental defects, particularly X-linked intellectual disability^28^. To date, the subcellular distribution of *O*-GlcNAcylated proteins (*O*-GlcNAcome) during the asymmetric stem cell divisions have not been investigated, owing primarily to the absence of appropriate high-resolution imaging probes.

Utilizing the *O*-GlcNAcase from *Clostridium perfringens* (*Cp*OGA), for which the crystal structure has been elucidated^29^, we have previously established hypo-*O*-GlcNAcylation *Drosophila* models^30^ and developed a tissue-specific *O*-GlcNAcylation profiling method^31,32^, revealing a fundamental connection between *O*-GlcNAcylation and cognitive function of the brain.

Here, we report the development of *O*-GlycoTracer, a live-cell imaging probe allows visualization of cellular *O*-GlcNAcome at high spatiotemporal resolution. The *O*-GlycoTracer unravels an unexpected asymmetric partition of *O*-GlcNAcome during the mitosis of *Drosophila* neuroblasts, which is largely mediated by 14-3-3 proteins. This unequal segregation of *O*-GlcNAcome is important for the binary cell fate determination, partially via creating structural and functional asymmetry in nuclear pores between the two daughters of neural stem cells. Our results reveal that the partition of cellular *O*-GlcNAcome is precisely orchestrated during asymmetric stem cell divisions, providing mechanistic insight into the etiology of *OGT* or *14-3-3* associated neurodevelopmental disorders.

## Results

### The O-GlcNAcome is asymmetrically segregated during neuroblast division

To visualize the spatiotemporal dynamics of the cellular *O*-GlcNAcome in living organisms, we developed the *O*-GlycoTracer by linking a fluorescent tag to a mutant OGA from *Clostridium perfringens* (D298N, *Cp*OGA^CD^) that is catalytically dead but retains a moderate binding activity toward the *O*-GlcNAc moiety^33^ (extended data fig. 1a). Transfection of *OGT* plasmid generated a palette of cells with varying cellular *O*-GlcNAcylation level. The bacterially purified *O*-GlycoTracer accurately labeled the cells with high *O*-GlcNAcylation level, comparable to the widely used anti-*O*-GlcNAc antibody RL2^34^ (extended data fig. 1b). The *O*-GlycoTracer also, as expected, detected a reduced level of *O*-GlcNAcylation in *Drosophila* embryos derived from an *Ogt* hypomorphic mutant^33^ (*Ogt/sxc^H^*^537^*^A^*) compared to those from wild type (extended data fig. 1c). Once injected or genetically expressed in early embryos, the *O*-GlycoTracer was enriched at the nuclear envelope, a region known to accommodate many *O*-GlcNAcylated substrates (extended data fig. 1d), and faithfully reported the decline in *O*-GlcNAcylation around the time of mid-blastula transition during *Drosophila* early embryogenesis^30^ (extended data fig. 1e). The control GFP-*Cp*OGA^DM^ (D298N, D401A) lacking the *O*-GlcNAc binding activity was diffusive in the embryonic cytoplasm, validating the specificity of the *O*-GlycoTracer in visualizing *O*-GlcNAcylation gradient in living cells.

*Drosophila* neuroblasts (NB) undergo asymmetric division, generating a self-renewing NB and a differentiating counterpart (fig. 1a). When the *O*-GlycoTracer was expressed in the NB lineage driven by *insc-Gal4*, it localized to interphase nucleus and nuclear envelope as observed in the early embryos (extended data fig. 1d). The signal then became diffusive in mitosis, and re-established the interphase pattern during mitotic exit (fig. 1b, supplementary video 1). Intriguingly, the *O*-GlycoTracer exhibited unequal distribution between the two daughter cells, with the future NB inheriting significant more signal in its nucleus (fig. 1c-1d). As a control, the GFP-*Cp*OGA^DM^ was evenly partitioned into the two daughter cells (extended data fig. 1f-1h, supplementary video 1). In subsequent symmetric divisions of ganglion mother cells (GMCs), the distribution of *O*-GlycoTracer appeared equal (fig. 1e-1g, supplementary video 2). These results indicate that the *O*-

**Figure 1.**
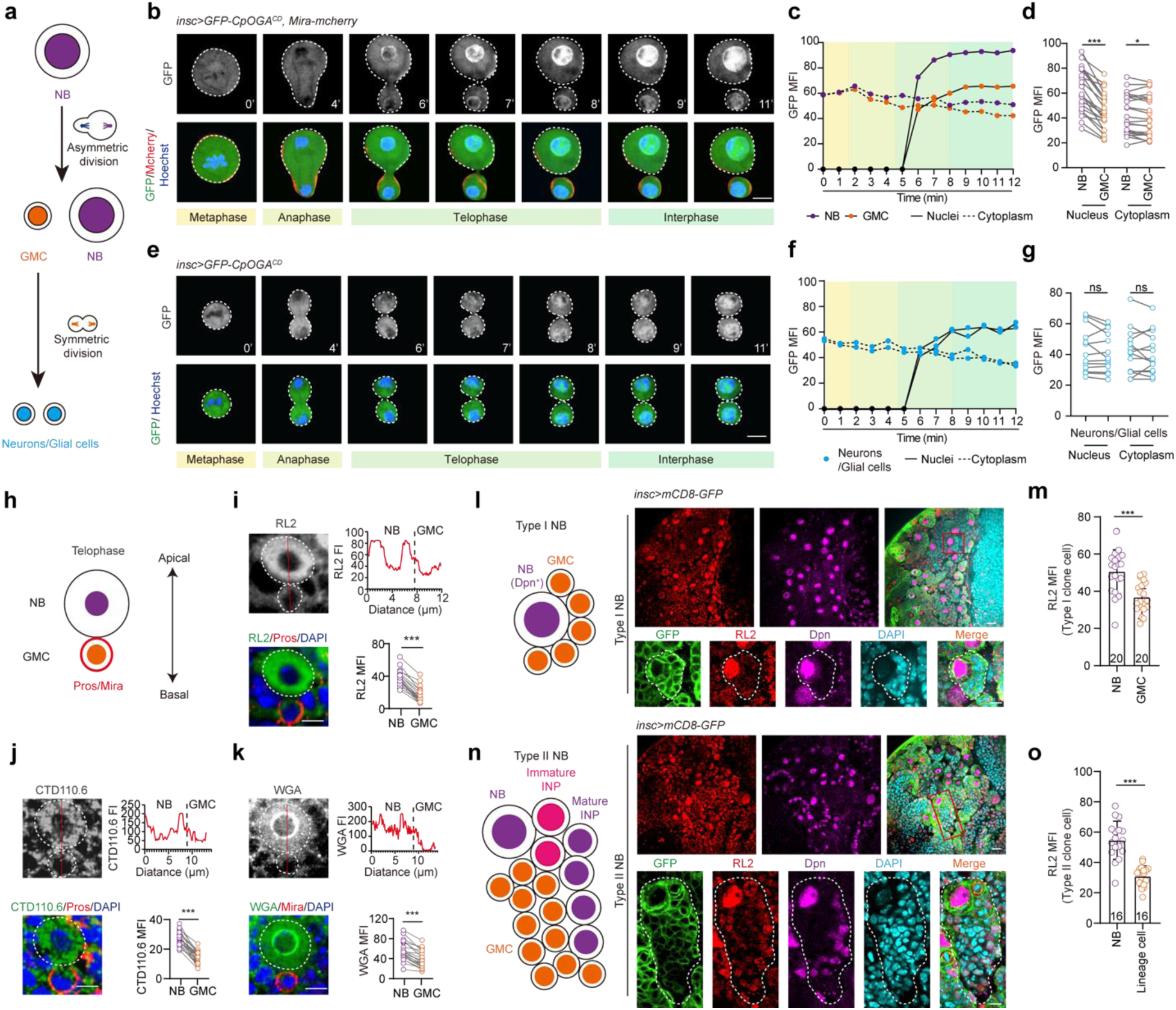
The *O*-GlcNAcome exhibits asymmetric segregation during neuroblast division. **a.** Diagram of *Drosophila* neuroblasts (NBs) divisions. NBs undergo asymmetric divisions to generate one NB and one ganglion mother cell (GMC). The GMCs further divide symmetrically to generate differentiated cells (neurons or glial cells). **b.** Real-time imaging of mitotic division of NB expressing *O*-GlycoTracer (GFP-*Cp*OGA^CD^) and Miranda (Mira)-mCherry. DNA is visualized with Hoechst (blue). NB and GMC are outlined by white dashed lines, with Mira-mCherry (red) labeling the GMC. Elapsed time is indicated in minutes. Scale bar, 5 μm. **c.** Quantification of the *O*-GlycoTracer mean fluorescence intensity (MFI) over time in the nucleus and cytoplasm of apical large cell (NB) and basal small cell (GMC) respectively. **d.** Quantification of nuclear and cytoplasmic *O*-GlycoTracer MFI in the two newly formed daughter cells at telophase (n = 22). **e.** Real-time imaging of symmetric division of GMC expressing *O*-GlycoTracer. DNA is visualized with Hoechst (blue). Cells are outlined by white dashed lines. Elapsed time is indicated in minutes. Scale bar, 5 μm. **f.** Quantification of the *O*-GlycoTracer MFI over time in the nucleus and cytoplasm of the two daughter cells of GMC. **g.** Quantification of nuclear and cytoplasmic *O*-GlycoTracer MFI in the two newly formed daughter cells of GMC at telophase (n = 13). **h.** Schematic of telophase of NB. The basal markers Mira and Prospero (Pros) localize to the GMC. **i-k.** Representative images of NB in telophase stained with anti-*O-*GlcNAc antibody RL2 (i), CTD110.6 (j), or lectin WGA (k). Mira or Pros is shown in red to label the GMC, and DNA is visualized with DAPI (blue). The fluorescence intensity (FI) plot of the staining along the red line is shown at the top right panel. Quantification result of MFI in NBs and GMCs is shown at the bottom right panel (n = 22). Scale bars, 5 μm. **l.** Diagram and representative images of Type I NB clone. Type I NBs produce GMCs that further divide again to form a pair of sibling neurons. NBs are marked by Deadpan (Dpn, magenta). *O*-GlcNAcylation is stained with anti-*O-*GlcNAc antibody RL2 (red). NBs and their lineage cells are marked with mCD8-GFP (green), and nuclei are visualized with DAPI (cyan). A Type I NB clone is outlined by white dashed lines. Scale bars, 10 μm (top row), 5 μm (bottom row). **m.** Quantification of RL2 MFI in NBs and GMCs within Type I NB clones (n = 20). **n.** Diagram and representative images of Type II NB clone. Type II NBs produce intermediate neural progenitors (INPs) which subsequently undergo multiple rounds of divisions to generate 4-6 GMCs. Both NBs and mature INPs are Dpn-positive (magenta). A Type II NB clone is outlined by white dashed lines. Scale bars, 10 μm (top row), 5 μm (bottom row). **o.** Quantification of RL2 MFI in NBs and their lineage cells (including INP and GMC) within Type II clones (n = 16). For statistical analyses, *P* values in (d), (g), (i), (j), and (k) are determined by two-tailed paired t test, while those in (m) and (o) are determined by two-tailed unpaired t test. The stars indicate significant differences (ns, not significant; \**p* < 0.05, \*\*\**p* < 0.001), error bars represent SD.

GlcNAcome is asymmetrically segregated during the mitosis of NBs. To confirm this observation made with the *O*-GlycoTracer, immunofluorescence staining of telophase NBs with anti-*O*-GlcNAc antibodies (RL2 and CTD110.6) or the plant-derived lectin wheat germ agglutinin (WGA) that recognizes *O*-GlcNAcylated substrates was performed (fig. 1h). Despite the difference in the subcellular pattern of fluorescent signal, likely caused by biased substrate preference or subtle asynchrony of the NBs at mitotic exit, the results unambiguously validated the preferential retention of *O*-GlcNAcome in the future neural stem cells (fig. 1i-1k). Conceivably, given the asymmetric allocation of *O*-GlcNAcome in mitosis, the NBs maintained the highest *O*-GlcNAcylation level among all the progeny cells within each NB clone (either Type I or Type II) in *Drosophila* larval brains (fig. 1l-1o).

### Optical screen reveals that 14-3-3 proteins mediate the unequal partition of O-GlcNAcome

To decipher the mechanism of action underlying the ordered segregation of *O*-GlcNAcome during NB division, we first examined mitotic partition of OGT/sxc and OGA, the two enzymes controlling *O*-GlcNAc cycling. Neither the endogenously GFP-tagged OGT/sxc nor OGA displayed uneven distribution between the two daughter cells at mitotic exit of NB (fig. 2a-2b, extended data fig. 2a), suggesting that the *O*-GlcNAc enzymes are not involved in generating the mitotic asymmetry of *O*-GlcNAcome. Previous biochemical studies have identified a list of proteins that are capable to bind *O*-GlcNAcylated substrates with their physiological functions left undetermined^35^. These candidate *O*-GlcNAc binders, according to the result of protein-protein interaction (PPI) analysis, are grouped into three functional modules (extended data fig. 2b, supplementary table 1). Of note, the module involved in the establishment of cell polarity contains 14-3-3ζ, 14-3-3ε, Moe, Chc, and tsr, many of which can interact with other known polarity factors governing the asymmetric division of NB^36,37^. We knocked down these *O*-GlcNAc binders that are highly expressed in NB by RNAi, and performed live-imaging screen using the *O*-GlycoTracer (fig. 2c-2d, extended data fig. 2c). Knockdown of 14-3-3ζ or 14-3-3ε, as well as tsr, significantly reduced the difference of *O*-GlycoTracer signal between the two NB progeny cells, indicating that they may mediate the asymmetric partition of *O*-GlcNAcome. Since both 14-3-3ε and 14-3-3ζ are contributing to the *O*-GlcNAcome asymmetry at mitotic exit, we examined their subcellular localization by immunofluorecence staining. The results revealed that 14-3-3 proteins themselves exhibited asymmetric distribution, preferentially localizing to the future neuronal stem cells (fig. 2e).

**Figure 2.**
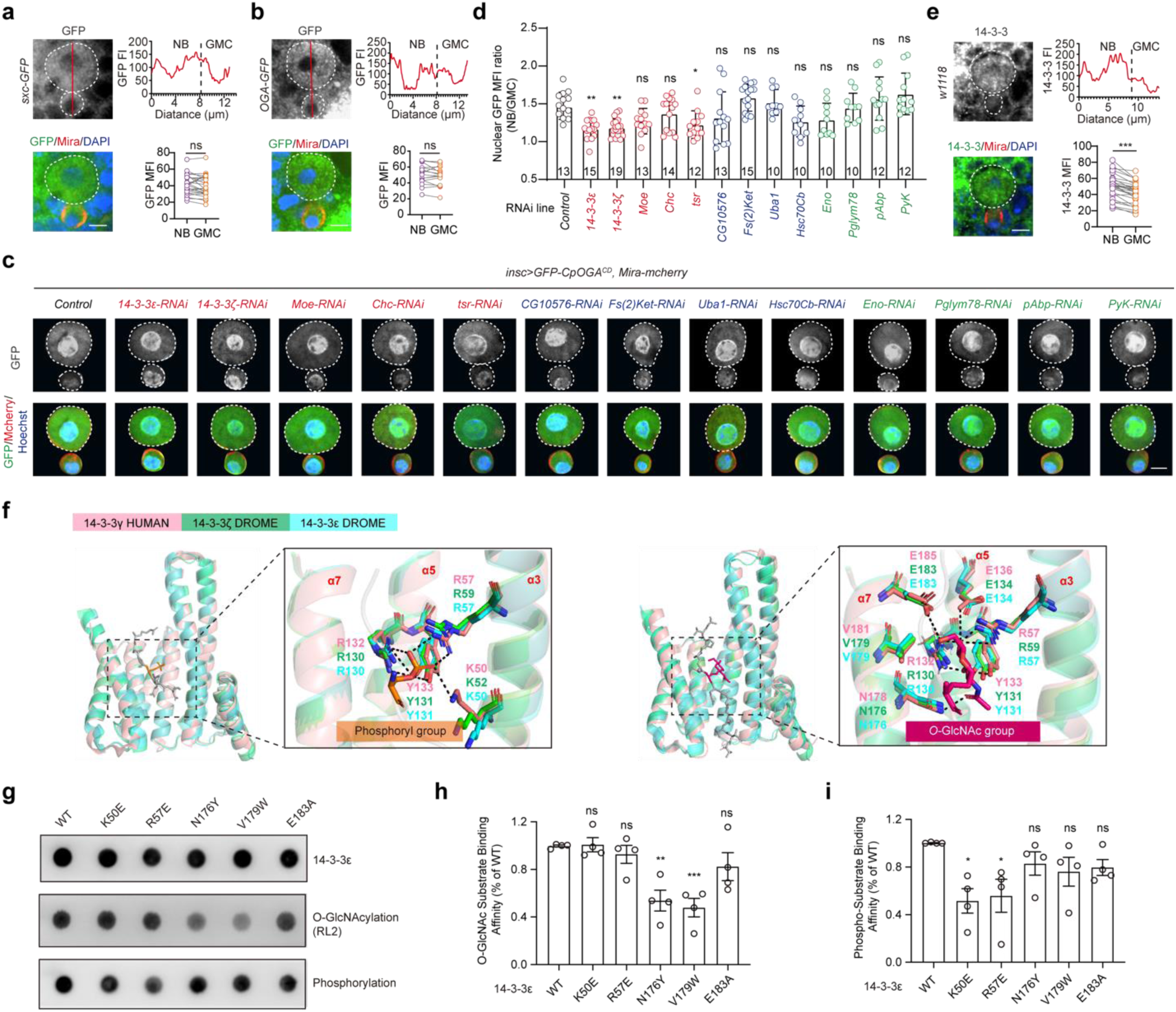
Candidate screen identifies 14-3-3 proteins as mediators of the unequal partition of *O-*GlcNAcome. **a-b.** Representative images of NBs at telophase expressing sxc-GFP (a) or OGA-GFP (b). Mira shown in red labels the GMC, and DNA is visualized with DAPI (blue). The fluorescence intensity (FI) plot of the GFP signal along the red line is shown at the top right panel. Quantification result of MFI in NBs and GMCs is shown at the bottom right panel (n = 25 and n = 17 respectively). Scale bars, 5 μm. **c.** Representative snapshots from live imaging of NBs at telophase. The expression of *O*-GlycoTracer and Mira-mCherry, along with shRNA targeting the indicated genes, are driven by *insc-Gal4*. DNA are visualized with Hoechst (blue) and Mira-mCherry (red) labels the GMCs. Scale bar, 5 μm. **d.** Quantification of the nuclear *O*-GlycoTracer MFI ratio of NBs over GMCs for the indicated genotypes (n = 10 - 19). **e.** Representative image of NB at telophase stained with anti-14-3-3 antibody (green). The fluorescence intensity (FI) plot of the staining signal along the red line is shown at the top right panel. Quantification result of MFI in NBs and GMCs is shown at the bottom right panel (n = 31). Scale bar, 5 μm. **f.** Structures of human 14-3-3γ (pink), *Drosophila* 14-3-3ζ (green) and *Drosophila* 14-3-3ε (cyan) bound to phosphopeptide (left, PDB 4J6S) or *O-*GlcNAc peptide (right, PDB 6BYJ). The key residues mediating the interaction are shown as sticks. The hydrogen bonds are visualized by black dashed lines. Specifically, the NHAc group at C2 position of *O*-GlcNAc forms hydrogen bond with N176 of *Drosophila* 14-3-3ε; the hydroxyl group at C3 position forms hydrogen bonds with R130 and Y131; the C4 hydroxyl group forms hydrogen bonds with R57, R130, and E134; the C5 hydroxyl group forms hydrogen bond with E183. **g.** Representative dot blot showing binding of purified wild-type or mutants *Drosophila* 14-3-3ε to substrates in lysates of adult flies. Membranes were probed with antibodies against 14-3-3ε, *O*-GlcNAc (RL2), and phosphoserine/threonine. **h-i.** Quantification of *O*-GlcNAc substrate (h) and phospho-substrate (i) binding capacities of wild-type and mutants *Drosophila* 14-3-3ε (n = 4). For statistical analyses, *P* values in (a), (b), and (e) are determined by two-tailed paired t test, while those in (d), (h), and (i) are determined by one-way ANOVA followed by Dunnett’s multiple comparison test. The stars indicate significant differences (ns, not significant; \**p* < 0.05, \*\**p* < 0.01, \*\*\**p* < 0.001), error bars represent SD.

Several human 14-3-3 isoforms are able to interact with *O*-GlcNAcylated substrates, and crystal structure demonstrates that human 14-3-3γ (YWHAG) recognizes *O*-GlcNAcylated as well as phosphorylated peptide via different residues^35^. Sequence analysis using ESPrit3.0 revealed a high degree of evolutionary conservation between *Drosophila* and human 14-3-3 proteins (extended data fig. 3, supplementary table 2). To dissect the molecular basis of 14-3-3 proteins in partitioning the *O*-GlcNAcome during mitosis of NB, we predicted the structures of the two 14-3-3 isoforms in *Drosophila* (14-3-3ζ and 14-3-3ε) using AlphaFold3, and performed pairwise structural alignment with human 14-3-3γ (fig. 2f). *Drosophila* and human 14-3-3 proteins consisted of nine antiparallel α-helices (α1-α9), four of which (α3, α5, α7, and α9) formed an amphipathic ligand-binding groove (fig. 2f, extended data fig. 3). Structural comparison indicated that the phosphate group of the phosphorylated peptide formed hydrogen bonds with residues K50, R57, R130, and Y131 of *Drosophila* 14-3-3ε, whereas the *O*-GlcNAc moiety formed hydrogen bonds with R57, R130, Y131, E134, N176, and E183 (PDB 4J6S and 6BYJ). Additionally, the *O*-GlcNAc moiety established hydrophobic interactions with V179 and the surrounding hydrophobic amino acids of *Drosophila* 14-3-3ε. To experimentally validate the recognition of *O*-GlcNAcylated substrates by *Drosophila* 14-3-3ε, we constructed *Drosophila* 14-3-3ε mutants that selectively disrupted the interaction with *O*-GlcNAc or phosphate group according to our structural analysis and previous report^35^, purified the wild-type and mutant 14-3-3ε proteins, and performed pulldown assays with lysates of adult flies (fig. 2g). Quantification of the dot blot results demonstrated that *Drosophila* 14-3-3ε could interact with both phosphorylated and *O*-GlcNAcylated proteins, and introduction of point mutations to the binding interface for phosphate (K50E, R57E) or *O*-GlcNAc (N176Y, V179W) attenuated the recognition of phosphorylated or *O*-GlcNAcylated substrates, respectively (fig. 2h-2i). Therefore, we reason that *Drosophila* 14-3-3 proteins function as *O*-GlcNAc binders, retaining the *O*-GlcNAcome preferentially into the future NB via their interactions with the polarity factors.

### Perturbation of the asymmetric distribution of O-GlcNAcome compromises neuronal cell fate determination and brain development

To evaluate the functional importance of the preferential retention of *O*-GlcNAcome in neuroblasts during brain development, we perturbed the *O*-GlcNAcylation homeostasis by ubiquitous expression of wild-type *Cp*OGA driven by *Da-Gal4*^30^ (*Cp*OGA^WT^). Third-instar larval brains expressing *Cp*OGA^WT^ or the control *Cp*OGA^DM^ were dissected and subject to single-cell RNA sequencing. Based on the expression of cell type-specific marker genes, four major cell types-NB lineage, neurons, glial cells, and trachea-were identified (fig. 3a, extended data fig. 4a-4b). Neurons in the mushroom body were annotated according to the expression of *prt* and *ey* as well. Cell-type composition analysis showed that the hypo-*O*-GlcNAcylation brains dissected from larva expressing *Cp*OGA^WT^ displayed a significant reduction in neurons, accompanied by a pronounced accumulation of cells in the NB lineage (fig. 3a). To refine the subpopulations in the NB lineage, re-clustering was performed, yielding six distinct groups according to the dynamic expression of marker genes associated with different stages of NB differentiation (fig. 3b, extended data fig. 4c-4d). Brains from the *Cp*OGA^WT^ group exhibited an increase in NBs (Type II) and intermediate neural progenitors (INPs), whereas the proportions of GMCs and new-born neurons were reduced compared to the control brains expressing *Cp*OGA^DM^ (fig. 3b). The shift in cell composition in the hypo-*O*-GlcNAcylation brains suggests that perturbation of *O*-GlcNAcylation homeostasis compromises cell fate determination during NB differentiation.

**Figure 3.**
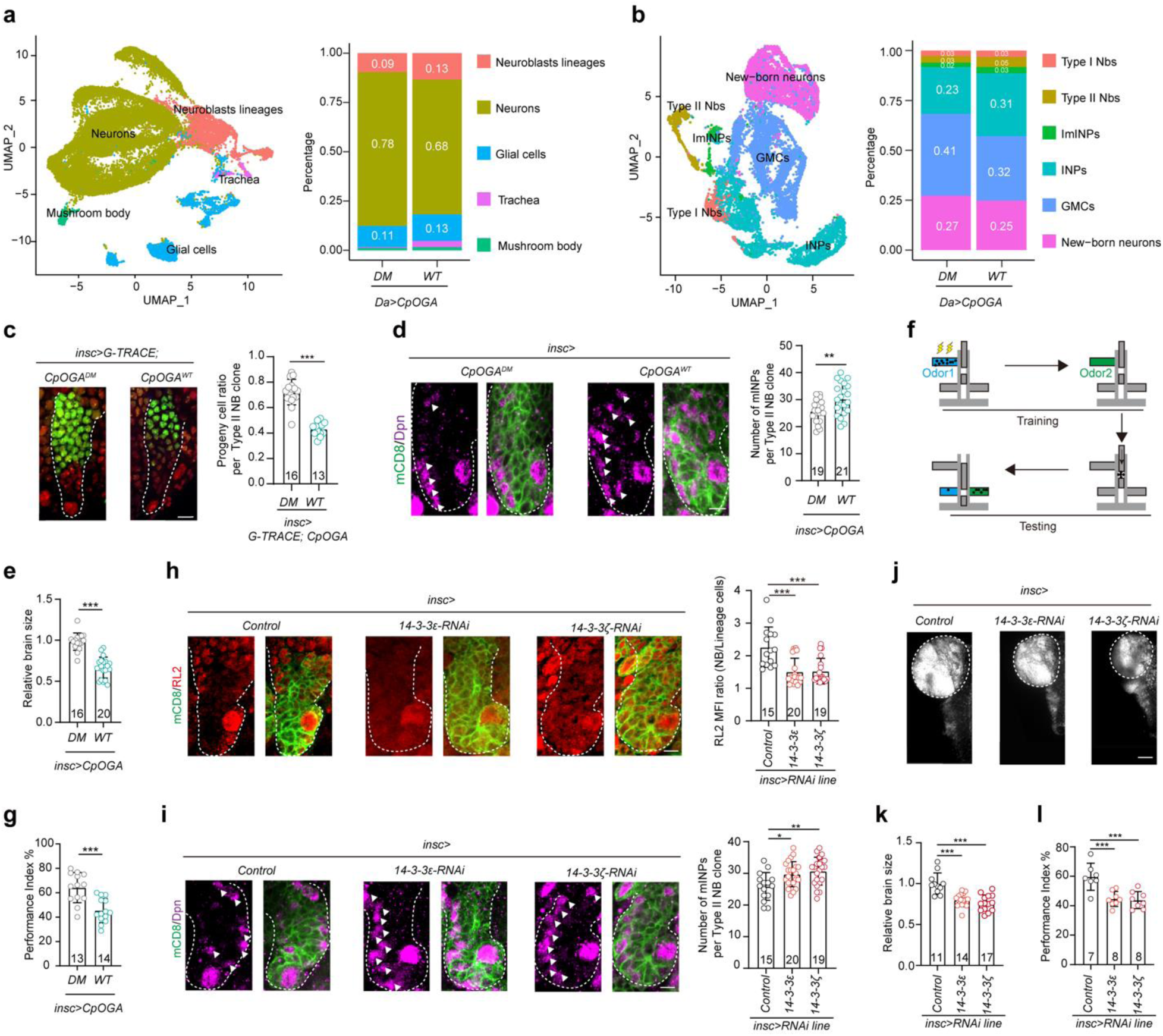
Interruption of the asymmetric allocation of *O-*GlcNAcome impairs binary cell fate decision of neuroblasts. **a.** Cell composition of the larval brains expressing the catalytically active *Cp*OGA^WT^ (WT) or its control *Cp*OGA^DM^ (DM). NB lineage was defined by the expression of *dpn*, *ase*, *wor* and *insc*; neurons were characterized by the expression of neurotransmitter-related genes such as *Syt4* and *brp*; glial cells were marked by the expression of *repo* and *wrapper*; trachea cells were defined by the expression of *grh* and *ImpE2*. Different cell types on a Seurat UMAP plot are color coded (left), and the stacked bar plot shows their percentage (right). **b.** Different subtypes of cells in the NBs lineage. Type I NBs were identified by the expression of *dpn*, *klu*, and *ase*; Type II NBs were characterized by the expression of *pnt*, *tll*, and *dpn*, with a notable absence of *ase*; Immature intermediate neural progenitors (imINPs) were defined by the expression of *pnt* and *erm*; mature INPs were distinguished by the expression of *dpn*, *ase*, *klu*, and *erm*; GMCs were marked by the presence of *ase* and absence of *dpn*; additionally, a population of new-born neurons was identified by the expression of *Hey*, *fne*, and *nSyb*. **c.** Representative images of cell lineage analysis for Type II NB clones expressing *Cp*OGA^WT^ or *Cp*OGA^DM^ using G-TRACE. The *insc-Gal4* drives the expression of RFP and FLP recombinase in NBs lineage (red). The FLP subsequently excises the STOP cassette flanked by FRT sites, enabling permanent expression of GFP in all daughter cells (green). The Type II NB clone is outlined by white dashed line. Scale bar, 10 μm. Quantification of the ratio of GFP-positive progeny cells in a clone is shown on the right (n = 16 and 13). **d.** Representative images of Type II NB clones expressing *Cp*OGA^WT^ or *Cp*OGA^DM^ stained with Dpn (magenta). The NBs and their progeny cells are marked with mCD8-GFP (green). Scale bar, 10 μm. Quantification of the number of Dpn-positive mature INPs per Type II NB clone is shown on the right (n = 19 and 21). **e.** Quantification of relative brain size at third instar larval stage of flies expressing *Cp*OGA^WT^ or *Cp*OGA^DM^ in the NB lineage (n = 16 and 20). **f.** Diagram of *Drosophila* learning assay. Approximately 100 flies were conditioned to link one of the two aversive odors (MCH or OCT) with an electric shock in the upper section of the T-maze. The flies were then moved to the lower section of the T-maze to test their learning ability by assessing the odor preference. **g.** A compilation of performance index of the adult flies expressing *Cp*OGA^WT^ or *Cp*OGA^DM^ in the learning test (n = 13 and 14). **h.** Representative images of Type II NB clones with control, 14-3-3ζ, or 14-3-3ε knockdown. *O*-GlcNAcylation is visualized with RL2 (red). The NBs and their lineage cells are marked with mCD8-GFP (green). The Type II NB clones are outlined by white dashed lines. Scale bar, 10 μm. Quantification of the ratio of RL2 MFI in NBs over their progeny cells is plotted on the right (n = 15 - 20). **i.** Representative images of Type II NB clones with control, 14-3-3ζ, or 14-3-3ε knockdown. NBs and mature INPs are labeled with Dpn (magenta). Cells within the NBs lineage are marked with mCD8-GFP (green). Scale bar, 10 μm. Quantification of the number of mature INPs per Type II NB clone is shown on the right (n = 15 - 20). **j.** Representative images of third instar larval brains with control, 14-3-3ζ, or 14-3-3ε knockdown in the NBs lineage. Nuclei are visualized by DAPI (grey). Brain lobes are outlined by white dashed lines. Scale bar, 50 μm. **k.** Quantification of relative brain size after knockdown of control, 14-3-3ζ, or 14-3-3ε. (n = 11 - 17). **l.** A compilation of performance index of the adult flies with control, 14-3-3ζ or 14-3-3ε knockdown in the learning test (n = 7 - 8). For statistical analyses, *P* values in (c), (d), (e), and (g) are determined by two-tailed unpaired t test, while those in (h), (i), (k), and (l) are determined by one-way ANOVA followed by Dunnett’s multiple comparison test. The stars indicate significant differences (\*\**p* < 0.01, \*\*\**p* < 0.001), error bars represent SD.

To delineate the differentiation defect observed in single-cell RNA sequencing analysis, we overexpressed *Cp*OGA^WT^ and the control *Cp*OGA^DM^ specifically in NB lineage cells using the *insc-Gal4* driver. Immunostaining with anti-*O*-GlcNAc antibody RL2 confirmed that overexpression of *Cp*OGA^WT^ reduced *O*-GlcNAcylation level in NB clones (extended data fig. 5a-5b), and diminished the difference between NBs and their progeny cells (extended data fig. 5c). Lineage tracing analysis using G-TRACE revealed a significant reduction of differentiated progeny cells in the Type II NB clones expressing *Cp*OGA^WT^ (fig. 3c). Accordingly, within these clones the number of INPs was increased (fig. 3d). These results were consistent with that from single-cell RNA sequencing analysis. Notably, the overall brain size of larva expressing *Cp*OGA^WT^ in the NB lineage was significantly smaller than that expressing the control *Cp*OGA^DM^ (fig. 3e). To determine whether this neurodevelopmental abnormality translates into functional deficit in adult flies, olfactory associative learning assay was performed as previously reported^31^ (fig. 3f). The results showed that flies expressing *Cp*OGA^WT^ in the NB lineage performed poorly in linking odor with electric shock compared to the *Cp*OGA^DM^ control (fig. 3g, extended data fig. 5d), suggesting an impairment of cognitive function.

To directly assess the importance of unequal partition of *O*-GlcNAcome during brain development, we repeated the cytological and functional analyses with 14-3-3 knockdown flies (fig. 2d, extended data fig. 2b). Knockdown of 14-3-3ε or 14-3-3ζ using the *insc-Gal4* compromised the asymmetric partition of *O*-GlcNAcome (fig. 2c). Accordingly, the difference of *O*-GlcNAcylation level between NBs and their progeny cells was reduced (fig. 3h). Moreover, similar to the observation in *Cp*OGA^WT^ expressing flies, the number of INPs within each Type II NB clones was increased in the 14-3-3 knockdown brains (fig. 3i). Furthermore, knockdown of 14-3-3ε or 14-3-3ζ resulted in smaller brains in the third-instar larva (fig. 3j-3k), and when tested with the olfactory learning assay, these flies also exhibited deficits in neuronal functions (fig. 3l).

The observed cognitive impairment of 14-3-3 knockdown flies is reminiscent of phenotypes of human neurodevelopmental disorders. Similar to that of *OGT/sxc*^28^, mutations in the *14-3-3* genes have been identified in patients with neurodevelopmental disabilities^38–40^. We curated these pathogenic mutations and mapped them onto the three-dimensional structure of 14-3-3 proteins (supplementary table 3). Most of the affected residues were highly conserved between human and *Drosophila*, and more importantly, they tended to locate within the substrate-binding pocket of 14-3-3 (extended data fig. 6). These results suggest that disruption the binding of 14-3-3 to its substrates, including *O*-GlcNAcylated proteins, may contribute to the pathology of neurodevelopmental disorders in humans.

### The nucleoporin Nup153 exhibits O-GlcNAcylation-dependent asymmetric segregation

To identify major *O*-GlcNAcylated substrates that are asymmetrically segregated during the mitotic division of NBs, we employed a previously described tissue-specific profiling method based on TurboID^31^ to capture putative *O*-GlcNAcylated proteins in NB lineage (extended data fig. 7a). After fed the larva with biotin-supplemented food, *O*-GlcNAcylated proteins in Type I and Type II NB clones were selectively biotinylated, allowing for streptavidin enrichment and mass spectrometry analysis (extended data fig. 7b-7c). A total of 624 putative *O*-GlcNAcylated substrates were identified (supplementary table 4). KEGG pathway analysis revealed that they were most significantly enriched in nucleocytoplasmic transport (dme03013), followed by valine, leucine and isoleucine degradation (dme00280), the citrate cycle (dme00020), proteasome (dme03050), etc (fig. 4a, supplementary table 5).

**Figure 4.**
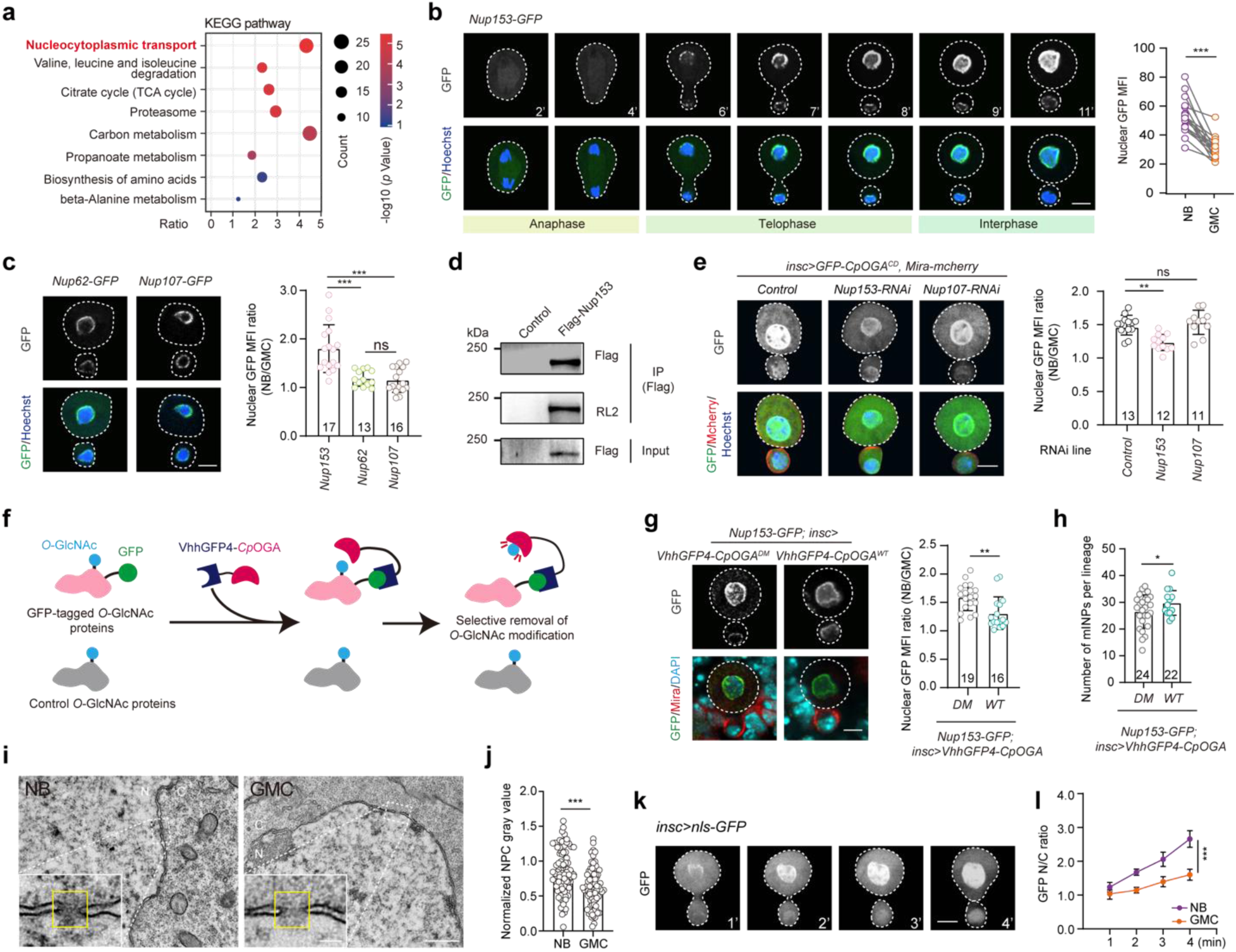
Nup153 is one of the key *O-*GlcNAcylated proteins that segregate asymmetrically. **a.** Gene ontology (GO) enrichment analysis of putative *O-*GlcNAcylated proteins detected in NB lineage. Bubble color indicates the -log10 (*p* value), and bubble size represents the ratio of genes in each category. **b.** Real-time imaging of asymmetric cell division of NBs from Nup153-GFP knockin flies. DNA is stained with Hoechst (blue). Elapsed time is indicated in minutes. NBs and GMCs are outlined by white dashed lines. Scale bar, 5 μm. Quantification of nuclear GFP MFI at telophase in the two daughter cells is shown on the right (n = 17). **c.** Representative snapshots from live imaging of NBs at telophase of asymmetric cell division from Nup62-GFP knockin or Nup107-GFP knockin flies. NBs and GMCs are outlined by white dashed lines. Scale bar, 5 μm. Quantification of the nuclear GFP MFI ratio of NBs and GMCs for the indicated genotypes is shown on the right (n = 13 - 17). **d.** Immunoprecipitation of Flag-tagged Nup153. *O-*GlcNAcylation of Nup153 is detected using anti-*O-*GlcNAc antibody RL2. **e.** Representative snapshots from live imaging of NBs at telophase after knockdown of the indicated genes. The *O*-GlcNAcome is visualized with the *O*-GlycoTracer (green), GMCs are labeled by Mira-mCherry (red), and DNA is stained with Hoechst (blue). NBs and GMCs are outlined by white dashed lines. Scale bar, 5 μm. Quantification of nuclear *O*-GlycoTracer signal in NBs over GMCs for the indicated genotypes is shown on the right (n = 11 - 13). **f.** Schematic of targeted de-*O-*GlcNAcylation strategy based on GFP nanobody (VhhGFP4) and catalytically active *Cp*OGA. **g.** Representative images of Nup153-GFP knockin neuroblasts at telophase expressing VhhGFP4-*Cp*OGA^WT^ or VhhGFP4-*Cp*OGA^DM^. The basal GMC marker Mira is shown in red. Scale bar, 5 μm. Quantification of the nuclear GFP MFI ratio of NBs over GMCs for the indicated genotypes is shown on the right (n = 19 and 16). **h.** Quantification of the number of mature INPs indicated by Dpn staining in each Type II NB clone of the indicated genotypes (n = 24 and 22). **i.** Representative TEM images showing mature NPCs (nuclear pore complex, indicated by yellow box) in NBs and the surrounding GMCs. N, nucleus; C, cytoplasm. Scale bars, 100 nm (left), 500 nm (right). **j.** Quantification of the normalized electronic density value within the nuclear pores (n = 84 and 76). **k.** Real-time imaging of NBs expressing nls-GFP during mitotic exit. The future NB and GMC are outlined by white dashed lines. Elapsed time is indicated in minutes. Scale bar: 5 μm. **l.** Quantification of relative nuclear GFP fluorescent intensity overtime in NB and GMC. (n = 4). For statistical analyses, *P* values in (b) are determined by two-tailed paired t test, while those in (g), (h), and (j) are determined by two-tailed unpaired t test. *P* values in (c) and (e) are determined by one-way ANOVA followed by Dunnett’s multiple comparison test. *P* value in (l) is determined by two-way ANOVA. The stars indicate significant differences (ns, not significant; \**p* < 0.05, \*\**p* < 0.01, \*\*\**p* < 0.001), error bars represent SD.

Taking into consideration that the *O*-GlycoTracer was heavily enriched on nuclear envelope, we compared the nuclear pore subunits identified in our analysis with those reported in previous studies^41^ (supplementary table 6), and selected components in the outer ring (Nup107), the central channel (Nup62), and the nuclear basket (Nup153) for endogenous GFP tagging (extended data fig. 8a). Live imaging showed that all the tagged nucleoporins were correctly reassembled into the nuclear envelope at mitotic exit of NBs. Intriguingly, while Nup107-GFP and Nup62-GFP were evenly distributed between the two daughter cells (extended data fig. 8b-8c), Nup153-GFP exhibited unequal segregation, with the daughter cell that maintains the stem cell fate inheriting a larger portion (fig. 4b-4c). This observation suggests that Nup153 may be one of the major *O*-GlcNAcylated proteins that are selectively retained in the future neural stem cells during NBs division.

Western blot analysis of immunoprecipitated Nup153 confirmed that it was modified by *O*-GlcNAcylation (fig. 4d). If Nup153 were the main *O*-GlcNAcylated substrates undergo asymmetric segregation, depletion of Nup153 should compromise the unequal distribution of the *O*-GlycoTracer signal. Indeed, when Nup153 was knocked down by RNAi in NBs, not only the overall intensity of *O*-GlycoTracer on nuclear envelope but also the difference of *O*-GlycoTracer signal between the two daughter cells was reduced (fig. 4e, extended data fig. 8d). To determine whether the biased segregation of Nup153 relies on *O*-GlcNAcylation, we designed a nanobody-based VhhGFP4-*Cp*OGA system that allows selective removal of *O*-GlcNAcylation from targeted substrates (fig. 4f). The effectiveness of this strategy was validated with immunoprecipitated Nup153 from S2 cells (extended data fig. 9a). When transgenically expressed in NBs of Nup153-GFP larva, VhhGFP4-*Cp*OGA^WT^ significantly decreased *O*-GlcNAcylation on nuclear envelope compared to the VhhGFP4-*Cp*OGA^DM^ control (extended data fig. 9b). Transmission electron microscopy (TEM) analysis of nuclear pores in the NBs showed a significant reduction in the electron density of the central channel in the VhhGFP4-*Cp*OGA^WT^ group (extended data fig. 9c). Moreover, live imaging of Nup153-GFP revealed that the presence of VhhGFP4-*Cp*OGA^WT^ disrupted the unequal distribution of Nup153-GFP at mitotic exit of NBs (fig. 4g). The number of INPs in the VhhGFP4-*Cp*OGA^WT^ expressing Type II NB clones was also increased (fig. 4h), indicative of compromised cell fate determination in NB lineage as seen in 14-3-3 knockdown brains. Based on these results, we conclude that *O*-GlcNAcylated Nup153 is a major *O*-GlcNAcylated substrates undergo asymmetric segregation during NBs division.

To further determine the structural and functional importance of the *O*-GlcNAcylation dependent asymmetric partition of Nup153, we performed TEM to compare the ultrastructure of nuclear pores in NBs and their progeny cells. In comparison to that of GMCs, the nuclear pores of NBs displayed higher electron density in the central channel (fig. 4i-4j). Accordingly, the nuclear importing of nls-GFP, which requires the nuclear pore activity, was significantly faster in NBs than in GMCs (fig. 4k-4l). Either decreasing the *O*-GlcNAcylation level by expressing *Cp*OGA^WT^, or disrupting the asymmetric allocation of *O*-GlcNAcome by knocking down of 14-3-3ε, resulted in decreased electron density in the central channel of nuclear pores in NBs (extended data fig. 9d-9e). Moreover, the difference in the nuclear importing rate of nls-GFP between the two daughter cells of NBs was markedly reduced (extended data fig. 9f-9g). These results imply that the asymmetric partition of the *O*-GlcNAcome during NBs division reported in this study may generate different nuclear pore activities between the two daughter cells, promoting the bifurcation of their cell fates.

## Discussion

Asymmetric inheritance of cellular components during mitosis plays a central role in balancing stem cell self-renewal with differentiation. This concept has been nourished by a growing body of observations across multiple model systems. Early cytological analysis of Ascidian embryos linked unequal distribution of cytoplasmic contents to the divergence of cell fates^42^. Decades later, genetic studies in *Drosophila* neuroblasts identified key fate determinants-such as Numb and Prospero-that are asymmetrically segregated during mitosis to influence lineage specification^2,3^. This fundamental principle has since been extended beyond fate determinants to include organelles: centrosomes in *Drosophila* male germline stem cells, mitochondria in stem-like human mammary epithelial cells, and lysosomes in hematopoietic stem cells all exhibit asymmetric partitioning during divisions^7,11,20^. Moreover, recent findings have revealed that even genetic and epigenetic materials are subject to asymmetric segregation-for instance, the biased segregation of the template sister chromatid of sex chromosomes and the preferential retention of old histone H3 in *Drosophila* male germline stem cells^14,16^. Additionally, in embryonic and young mouse neural stem cells, damaged proteins tagged with ubiquitin are selectively inherited by the differentiating daughter cell, a process that diminishes with age^19^. Here, we offer a post-translational modification-elicited proteome-wide view of asymmetric inheritance, particularly with respect to *O*-GlcNAcylation which is essential for the maintenance of stemness. With our *O*-GlycoTracer, we discovered that the *O*-GlcNAcome is unequally partitioned during *Drosophila* neuroblast division. This finding reveals a previously unrecognized layer of cellular asymmetry, demonstrating that the metabolism-linked post-translational modifications can be inherited in a biased manner. We propose that the proteome-wide asymmetry during mitosis serves as a regulatory mechanism to ensure binary cell fate determination of the two daughters, providing fresh insight into how stem cells integrate intrinsic metabolic state to direct lineage outcomes.

At the mechanistic level, our study identified 14-3-3 proteins as major mediators of the asymmetric partition of *O*-GlcNAcome during neuroblast division. 14-3-3 proteins have been reported to recognize the saccharide moiety of *O*-GlcNAcylated substrates and proposed to be *O*-GlcNAc “reader” to effect downstream signaling^35^. Yet, the binding affinity between 14-3-3 and glycopeptide is low, leaving its physiological function in doubt. Instead of “reader”, we regard 14-3-3 proteins as “sorter” of *O*-GlcNAcylated targets during neural stem cell division, probably through transient and multivalent engagements with the cellular *O*-GlcNAcome. The sorting process during neuroblast division may involve their interactions with the polarity factors. The neuroblasts establish transient apical-basal polarity during mitosis^36^, with the apical cortex defined by actin-dependent recruitment of atypical protein kinase C (aPKC), which interacts with Bazooka (Baz/Par-3) and Par-6 to form the evolutionarily conserved Par complex^43,44^. Apical polarity directs mitotic spindle orientation through Inscuteable, which links the spindle-associated Pins–Gαi–Mud complex to Baz/Par-3, aligning the spindle along the polarity axis^45–48^. Meanwhile, aPKC activity restricts cell fate determinants such as Miranda, Prospero, Brat, and Numb to the basal cortex^4,49–51^, priming the basal daughter cell for differentiation. 14-3-3, also known as Par-5, is an evolutionarily conserved regulator of cell polarity. In *Drosophila*, 14-3-3 proteins interact with Par-1 to regulate the polarization of the anterior-posterior (A-P) axis during oogenesis^52^. 14-3-3 proteins can also bind Baz/Par-3 in a phosphorylation dependent manner, concentrating the Par complex to the apical membrane^53^. We propose that 14-3-3 proteins via the interactions with the polarity machinery retain the *O*-GlcNAcome preferentially in the apical daughter cell maintaining stem cell identity. Of note, the interactions of 14-3-3 proteins with the Par complex as well as the *O*-GlcNAc moiety are conserved in mammals^54,55^, warranting future validations of their *O*-GlcNAc “sorter” activity in other model systems.

Among the thousands of proteins that can be *O*-GlcNAcylated, our results suggested that nuclear pore complex, particularly Nup153 in the nuclear basket, is one of the major *O*-GlcNAcylated cargos subject to the 14-3-3 mediated asymmetric partition during neuroblast division. Interestingly, mammalian neural stem cells also have the highest level of Nup153 compared to their progeny^56^. The 14-3-3 mediated and *O*-GlcNAcylation dependent sorting of nuclear pore activities might be vital for the proper development of brain. Remarkably, human genetic mutations at multiple nodes in this process have been identified in patients with various neurodevelopmental disorders (NDDs). Many disease-associated variants have been mapped to the substrate-binding TPR domain as well as the catalytic region of human *OGT* in patients suffering from an X-linked intellectual disability syndrome. Similarly, mutations of human *14-3-3*, particularly those around the cargo-binding pocket, are associated with a spectrum of neurodevelopmental abnormalities. For example, human *YWHAG/14-3-3γ* mutations have been recognized as a genetic cause of developmental and epileptic encephalopathies^39,40^ (DEEs), and *YWHAZ/14-3-3ζ* mutations have been linked to intellectual disability (ID), global developmental delay (GDD), and cardiofaciocutaneous syndrome^38^ (CFCS). Besides, mutations in nucleoporins are linked to cell type-specific NDDs: Nup62 mutations lead to infantile bilateral striatal necrosis^57^, Nup214 mutations cause progressive encephalopathy with cortical atrophy ^58,59^, nuclear basket protein TPR variants result in severe intellectual disability^60^, and homozygous Nup107 mutations cause microcephaly with steroid-resistant nephrotic syndrome^61^. Of particular interest, patients suffering from these differernt NDDs often share overlapping synptoms including neurodevelopmental delay, microcephaly, and intellectual disability, which was also observed in our *Drosophila* models, suggesting that our finding in *Drosophila* is relevant to the cellular mechanisms underpinning these neurodevelopmental disorders in human.

In conclusion, our work reveals a regulatory axis in which 14-3-3 dependent *O*-GlcNAcome sorting orchestrates the asymmetric inheritance of cellualr proteins, establishing proteome-level asymmetry as a previously unrecognized layer of stem cell regulation. This study also provides a conceptual framework linking *O*-GlcNAc homeostasis to neurodevelopmental integrity and offers mechanistic insight into how mutations in *OGT*, *14-3-3*, and nucleoporin genes may disrupt the development of the brain.

## Materials and Methods

### Drosophila Stocks and Genetics

All *Drosophila* stocks were maintained on standard cornmeal medium at 25°C under a 12-h light/dark cycle. The strains used included: *w*^1118^, *;Sco/CyO;TM3/TM6B*, *;;Da-Gal4*, *;insc-Gal4;*, *;;UAS-mCD8-GFP* (gift from Zhuohua Zhang), *;;mira-HA-mCherry/TM3* (gift from Sijun Zhu), *;sxc^H^*^537^*^A^;*(gift from Daan M.F. van Aalten), *;MTD-Gal4;* (BDSC #31777), *;Pros-GFP;* (BDSC #66463), *;;EGFP-OGA* (BDSC #91771), *;GFP-Nup107;* (BDSC #35514), *;UAS-Nls-GFP;* (BDSC #4775), *G-TRACE* (gift from Hai Huang), *;;UAS-HA-TurboID-CpOGA^CD^* and *;;UAS-HA-TurboID-CpOGA^DM^*. RNAi lines were *;UAS-shPros;* (VDRC #v330051), *;UAS-shMyc;* (BDSC #2947/2948), *;;UAS-sh14-3-3ε* (THU4849, THU1467), *;;UAS-sh14-3-3ζ* (THU2964, THU4850), *;;UAS-shMoe* (THU0542), *;;UAS-shChc* (THU1462, THU2693), *;;UAS-shtsr* (THU0972), *;;UAS-shCG10576* (THU3722), *;;UAS-shFs(2)Ket* (THU2731, THU5823), *;;UAS-shUba1* (THU2127), *;;UAS-shHsc70Cb* (THU3637), *;UAS-shEno;* (TH201500405.S, TH02508.N), *;;UAS-shPglym78* (THU2186, TH02536.N), *;UAS-shpAbp;* (TH02923.N, THU0967), *;UAS-shPyk;* (TH03685.N), *;UAS-shNup107;* (TH04231.N), and *;;UAS-shwhite* (THU0558). Transgenic lines for *;;UAS-GFP-CpOGA^WT^*, *;;UAS-GFP-CpOGA^CD^*, *;;UAS-GFP-CpOGA^DM^*, *;;UAS-HA-CpOGA^WT^*, *;;UAS-HA-CpOGA^DM^*, *;;UAS-HA-VhhGFP4-CpOGA^WT^*, and *;;UAS-HA-VhhGFP4-CpOGA^DM^*were generated by cloning the respective coding sequences into the pUASz vector via Gibson Assembly (TransGen, CU201-02). The resulting attB-containing constructs were injected into *y1, w67c23; P(CaryP)attP2* embryos for φC31 integrase-mediated site-specific integration, and stable F1 transgenic lines were established by crossing G0 adults to double balancer stocks.

### Endogenous knock-in lines via CRISPR/Cas9

Endogenous EGFP-tagged fly lines for *Nup153*, *Nup62*, and *sxc* were generated using the CRISPR/Cas9 system. Guide RNAs (gRNAs) were designed to target the intended genomic loci using the flyCRISPR target finder tool (https://flycrispr.org/) based on sequences from FlyBase. The gRNAs were cloned into the pU6-BbsI-chiRNA vector (Addgene #45946) and co-injected with donor plasmids carrying the EGFP tag (T-EGFP) into embryos of the *vasa-Cas9* strain (BDSC 51324/78781); all injections were performed by UniHuaii Co., Ltd (Zhuhai, China). For *Nup62* and *sxc*, F0 adults were crossed with double balancer flies (*Sco/CyO; TM3/TM6B*); F1 males carrying the *CyO* balancer were then mated with double balancer females, and their offspring were genotyped by PCR. For *Nup153*, F0 adults were crossed with *FM6* flies (*FM6/FM6*), and F1 females carrying the *FM6* balancer were mated with *FM6* males (*FM6/Y*) before PCR genotyping. All edited loci were confirmed by Sanger sequencing. The resulting EGFP-tagged strains were maintained at 25°C, and the gRNA sequences and plasmid details are provided in supplementary table 7.

### Cell cultures

HeLa cells (Meisen, CTCC-0306) were maintained in high-glucose DMEM (Biological Industries, 01-052-1A) supplemented with 10% FBS (Meisen, CTCC-002-001) at 37°C under a humidified atmosphere of 5% CO₂. Transfection of plasmids encoding GFP-tagged human OGT was performed using Lipofectamine 2000 (Invitrogen, L3000015) according to the manufacturer’s instructions. *Drosophila* S2 cells were cultured in Schneider’s Insect Medium (Gibco, 21720024) containing 10% FBS at 25°C. Transfection of S2 cells with plasmids expressing Flag-Nup153, Flag-GFP-Nup153, HA-VhhGFP4-*Cp*OGA^WT^, or HA-VhhGFP4-*Cp*OGA^DM^ was carried out using Effectene Transfection Reagent (Qiagen, 301425) following the manufacturer’s protocol.

### Protein purification and fluorescent labeling

The cDNA encoding a truncated *Cp*OGA (amino acids 31-618) and its derivative mutants were cloned into a pET28 vector, generating constructs with an N-terminal HaloTag and a C-terminal 6×His-tag. Similarly, cDNA for full-length *Drosophila* 14-3-3ε wild-type and its mutants, as well as PCNA, were inserted into the pET28 vector, yielding N-terminal GFP- and mCherry-tagged proteins, respectively, each with a C-terminal 6×His-tag. All constructs were transformed into *Escherichia coli* Rosetta (DE3) cells. Transformed cells were first cultured overnight at 37°C in 3 mL of Luria-Bertani medium supplemented with 50 µg/mL kanamycin. This pre-culture was then diluted into fresh medium at 1:1000 and grown until the OD600 reached 0.6. Protein expression was induced by adding 0.5 mM isopropyl β-D-1-thiogalactopyranoside (IPTG), followed by incubation at 16°C for 18 h. Cells were harvested by centrifugation at 4,000 × g for 15 min at 4°C. The pellets were resuspended in ice-cold lysis buffer (20 mM Tris, 500 mM NaCl, 10 mM imidazole, pH 8.0) containing protease inhibitors (0.2 mM PMSF and 5 µM leupeptin). Cell lysis was performed using a high-pressure homogenizer at 800 bar for three cycles. The lysate was clarified by centrifugation at 12,000 × g for 20 min at 4°C, and the supernatant was incubated with pre-equilibrated Ni-NTA agarose resin at 4°C for 2 h. The resin was washed three times with wash buffer (20 mM Tris, 500 mM NaCl, 20 mM imidazole, pH 7.9), and the bound proteins were eluted with elution buffer (20 mM Tris, 200 mM NaCl, 300 mM imidazole, pH 7.9). The eluates were dialyzed against a storage buffer containing 40 mM HEPES (pH 7.4) and 150 mM KCl. For HaloTag-*Cp*OGA proteins, labeling was achieved by incubating with an equimolar amount of HaloTag Alexa Fluor 488 or 660 ligands at room temperature for 30 min. Unconjugated dyes were removed by size-exclusion chromatography using a G-50 column. All purified proteins were aliquoted, flash-frozen in liquid nitrogen, and stored at -80°C.

### Pulldown assay

Adult *Drosophila* flies (n = 500) were collected and lysed in a buffer containing 2% SDS, 10% glycerol, and 62.5 mM Tris-HCl (pH 6.8), supplemented with a protease inhibitor cocktail (Sigma, P8340), 1 mM PMSF (Sigma, P7626), and 50 µM Thiamet-G (Selleck, S7213). The lysates were rotated at 4°C for 1 h and clarified by centrifugation at 13,000 × g for 30 min at 4°C. The total protein concentration was determined using a BCA assay kit (Beyotime, P0009), yielding approximately 10 mg/mL. For each pulldown reaction, 20 µg of purified 6×His-GFP-14-3-3ε (the bait) was pre-incubated with 2 mg of fly lysate protein overnight at 4°C with gentle rotation. Anti-His magnetic beads (BeyoMag, P2135) were prepared by washing twice with 1× PBS using a magnetic rack. The pre-incubated bait-lysate mixture was then added to 20 µL of the equilibrated bead slurry and incubated at 4°C for 2 h with rotation. After incubation, the beads were collected on the magnetic rack and washed three times with 500 µL of 1× PBS. Finally, the bound proteins were eluted by resuspending the beads in 100 µL of 1× SDS sample buffer containing 2% β-mercaptoethanol, followed by boiling at 95°C for 10 min.

### Dot-blot assay

For the dot-blot assay, 1.5 µL of each eluate was directly spotted onto a nitrocellulose (NC) membrane (5.5 × 1.5 cm, spot spacing 0.7 cm; Beyotime, FFN08). The membrane was air-dried at room temperature for 1 min. Subsequently, it was blocked with 5% BSA in PBST for 1 h at room temperature and then incubated with primary antibodies diluted in the blocking solution overnight at 4°C. The primary antibodies used were: rabbit anti-14-3-3ε (Diagbio, db13861; 1:1,000), mouse anti-*O*-GlcNAc (RL2, Abcam, ab2739; 1:1,000), and mouse anti-phosphoserine/threonine (BD Biosciences, 612549; 1:1,000). Following incubation, the membrane was washed three times for 5 min each with PBST. It was then probed with appropriate HRP-conjugated secondary antibodies (Thermo Fisher Scientific; 1:10,000) for 1 h at room temperature. After another series of three 5-min PBST washes, the signals were visualized using an ECL substrate (Millipore, WBKLS0050).

### Embryo microinjection and live imaging

Embryo microinjection was performed as previously described^62^. Briefly, embryos were manually dechorionated, aligned on a glass coverslip, and secured with double-sided tape. After desiccation for 4-6 min, they were covered with a 1:1 mixture of halocarbon oils 27 and 700 (Sigma, H8773 and H8898). Microinjection was carried out at nuclear cycles 9–10 using a manual micromanipulator (WPI) and a pneumatic picopump (WPI, SYS-PV830). The injection mixture contained Alexa Fluor 488-labeled HaloTag-*Cp*OGA proteins (10 µg/µL), PCNA-mCherry (2 µg/µL), and wheat germ agglutinin (WGA; 1.25 µg/µL; Thermo Fisher Scientific, W21404). Live imaging of the injected embryos was performed at room temperature on a ZEISS LSM 880 confocal microscope using a 63× Plan-Apochromat/1.4 NA oil immersion objective.

### Neuroblast live imaging

The imaging medium, consisting of Schneider’s insect medium supplemented with 10% FBS and

2.5 μg/mL Hoechst (Beyotime, C1022), was equilibrated to room temperature before use. Larvae were dissected at 72 h after hatching in the imaging medium. The isolated brains were then transferred to a stainless-steel slide and positioned within a cavity formed by a pre-attached 40 × 22 mm coverslip. Each brain was surrounded with approximately 7 µL of imaging medium and protected with a 20 × 20 mm coverslip. Time-lapse imaging of neuroblasts was performed at room temperature using a Nikon CSU-W1 spinning disk confocal system equipped with a 100× Plan-Apochromat 1.4 NA oil-immersion objective. Z-stacks spanning 8 μm with a step size of 1 μm were acquired at 1-min intervals.

### Odor avoidance and olfactory learning test

Behavioral assays were performed at 25℃ in an environmental chamber maintained at 70% humidity. To assess odor avoidance, flies were placed in the center of a T-maze apparatus and given a 2-min choice between air and an aversive odor, either 4-methylcyclohexanol (MCH; Sigma, 104191) or 3-octanol (OCT; Sigma, 218405). At the end of the assay, flies trapped in each arm were anesthetized and counted. The odor avoidance performance index (PIodor) was calculated as:

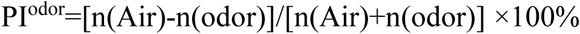

Flies exhibiting normal odor avoidance were subjected to olfactory learning tests. During training, flies were exposed to a conditioned odor (CS⁺) paired with an electric shock (60 V) for 1 min, followed by air, and then to a second odor (CS⁻) without shock. Flies were then tested in the T-maze, where they chose between CS⁺ and CS⁻ for 2 min. The learning performance index (PI) was calculated as:

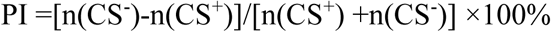

Each experiment consisted of two reciprocal trials, alternating MCH and OCT as CS⁺, and the mean PI was used to correct for odor preference bias. Each data point represented ∼200 flies (1:1 male-to-female ratio), with balanced training for both odors.

### Western blot assay

*Drosophila* S2 cells or fly tissues were lysed in a buffer containing 2% SDS, 10% glycerol, and 62.5 mM Tris-HCl (pH 6.8), supplemented with a protease inhibitor cocktail (Sigma, P8340, 1:100), 1 mM PMSF (Sigma, P7626), and 50 µM Thiamet-G (Selleck, S7213). The lysates were clarified by centrifugation at 13,000 × g for 30 min, and the supernatant protein concentration was quantified using a BCA assay kit (Beyotime, P0009). Protein samples were denatured by mixing with an equal volume of 2× SDS sample buffer containing 4% β-mercaptoethanol and boiling at 95°C for 10 min. The denatured proteins were separated on 10% SDS-polyacrylamide gels by electrophoresis (90 V for 30 min followed by 120 V for 60 min) and subsequently transferred onto PVDF membranes (Millipore, IPVH00010) at 290 mA for 90 min. The membranes were blocked with 5% skim milk for 1 h at room temperature and then incubated with specific primary antibodies diluted in 5% BSA (Biofroxx, 4240GR005) overnight at 4°C. The primary antibodies used were: streptavidin-HRP (GenScript, M00091, 1:2,000), anti-*O*-GlcNAc (RL2, Abcam, ab2739, 1:2,000), anti-HA (Cell Signaling Technology, 3724, 1:3,000), anti-Tubulin (Cell Signaling Technology, 12351S, 1:3,000), anti-FLAG (Cell Signaling Technology, 14793, 1:3,000), and anti-GFP (Abcam, ab290, 1:1,000). After incubation with the primary antibodies, the membranes were probed with appropriate HRP-conjugated secondary antibodies (Thermo Fisher Scientific, 1:10,000) for 1 h at room temperature. The immunoreactive signals were finally detected using an ECL substrate (Millipore, WBKLS0050).

### Immunoprecipitation

Third-instar larvae or S2 cells were lysed in RIPA buffer (50 mM Tris-HCl pH 8.0, 150 mM NaCl, 0.1% SDS, 0.5% sodium deoxycholate, 1% NP-40, 10 mM NaF, 10 mM Na₂VO₄, 50 µM Thiamet-G) supplemented with a protease inhibitor cocktail (Sigma, P8340, 1:100) and 1 mM PMSF (Sigma, P7626) on ice for 30 min. The lysates were clarified by centrifugation at 13,000 × g for 30 min at 4°C, and the protein concentration of the supernatants was determined using a BCA assay kit (Beyotime, P0009). For the immunoprecipitation, pre-washed Streptavidin magnetic beads (MCE, HY-K0208) or Anti-FLAG M2 Affinity Gel (Sigma, A2220) were incubated with equal amounts of the clarified lysates overnight at 4°C with constant rotation. Subsequently, the beads were collected and washed twice with 1 mL of RIPA buffer. The bound proteins were eluted by resuspending the beads in 1× SDS sample buffer and boiling at 95°C for 10 min. The eluted samples were stored at -80°C for subsequent western blot analysis.

### Protein identification by LC-MS/MS

Biotin food was prepared by adding 200 mM biotin (Merck, B4501) to hot (∼60 ℃) standard fly food and dissolved to a final concentration of 100 μM. *insc-Gal4* flies were crossed with *UAS-HA-TurboID-CpOGA^CD^* or *UAS-HA-TurboID-CpOGA^CD^* flies, and progeny were raised on biotin food until the third-instar larval stage. Larval brains (∼300 per sample) expressing TurboID-*Cp*OGA^CD/DM^ in neuroblast lineages were collected and subjected to streptavidin-based immunoprecipitation as described above. Bound proteins were separated by SDS-PAGE and lightly stained with Coomassie brilliant blue (Solarbio, C8430-10g). Gel slices were reduced with 10 mM DTT and alkylated with 55 mM IAA (Merck, I6125) prior to digestion.

LC-MS/MS was performed as previously described^31^. Raw data were processed in MaxQuant for protein identification and label-free quantification (LFQ). Perseus software was used to remove contaminants, reverse hits, and proteins identified only by site. A pseudo-count of 1 was added to protein intensities before log₂ transformation. Fold changes (log₂ FC) were calculated for TurboID-*Cp*OGA^CD^ samples relative to controls across three replicates. Only proteins identified with ≥2 unique peptides and present in at least two of three replicates were retained for further analysis. Statistical significance was determined using a two-tailed unpaired t-test. Proteins were considered putative *O*-GlcNAc substrates if they showed log₂ FC > 1 or *p* < 0.05.

### Immunostaining and image analysis

For immunostaining, fly tissues were fixed in 4% paraformaldehyde (PFA; Biosharp, BL539A) for 30 min at room temperature. After fixation, samples were washed three times with PBS (Biological Industries, 02-023-1A) and then permeabilized and blocked in 0.3% PBTN buffer (PBS containing 0.3% Tween-20 and 5% normal goat serum (NGS; Solarbio, SL038)) for 90 min at room temperature. The tissues were incubated with primary antibodies diluted in 0.3% PBTN overnight at 4°C, washed three times for 10 min each with 0.3% PBST (PBS with 0.3% Tween-20), and subsequently incubated with fluorophore-conjugated secondary antibodies (Thermo Fisher Scientific, 1:500) and DAPI (Sigma, D9542, 1:500) for 1 h at room temperature. Following three final washes with 0.3% PBST, samples were mounted in SlowFade Diamond Antifade Mountant (Invitrogen, S36963) for imaging. The primary antibodies and probes used were: chicken anti-GFP (Aves Labs, GFP-1010, 1:200), mouse anti-O-GlcNAc (RL2, Abcam, ab2739, 1:200), mouse anti-*O*-GlcNAc (CTD110.6, Sigma, O7764, 1:200), mouse anti-Miranda (gift from Dr. Seung Kim, 1:200), rat anti-Deadpan (Abcam, ab195173, 1:500), mouse anti-phosphoserine/threonine (BD Biosciences, 612549, 1:200), mouse anti-YWHAE (14-3-3ε, Diagbio, db13861, 1:200), Streptavidin-Cy3 (BioLegend, 405215, 1:200), and wheat germ agglutinin (WGA, Thermo Fisher Scientific, W21404, 1.25 µg/µL).

For embryo and HeLa cell immunostaining, established protocols were followed^30^, using Alexa Fluor 488-labeled HaloTag-*Cp*OGA (10 µg/µL) for embryos, and the RL2 antibody (1:400) or Alexa Fluor 660-labeled HaloTag-*Cp*OGA^CD^ (10 µg/µL) for HeLa cells. Images were acquired using a ZEISS LSM 880 or a Nikon CSU-W1 spinning disk confocal microscope. Quantitative image analysis, including measurements of fluorescence intensity, area, particle count, and size, was performed using ZEN (Zeiss), NIS-Elements (Nikon), and ImageJ software.

### RT-qPCR

Total RNA was extracted from fly samples using TRIzol reagent (Thermo Fisher Scientific, 15596026). Subsequently, 1 µg of total RNA was reverse-transcribed to generate cDNA using the RevertAid First Strand cDNA Synthesis Kit (Thermo Fisher Scientific, K1621). Quantitative PCR (qPCR) was then performed using the SYBR Green qPCR Master Mix (Solomon Biotech, QST-100) on a QuantStudio 3 Real-Time PCR System (Applied Biosystems). The reaction mixtures were prepared with the synthesized cDNA as the template. The ribosomal gene *RpL32* was used as the internal reference for normalization, and the relative expression levels of target genes were calculated using the comparative ΔΔCt method. All primer sequences are provided in supplementary table 7.

### Brain dissection, dissociation and single-cell suspension preparation

Third-instar larvae (100-150) were briefly rinsed with water to remove food debris and transferred to a petri dish lid containing drops of ice-cold PBS. Brains were dissected using fine forceps, and after removal of the ventral nerve cord, the intact brain lobes were collected in a low-DNA-binding tube containing 250 μL of PBS. The samples were then processed for single-cell suspension preparation. Briefly, tissues were pelleted by centrifugation at 500 × g for 5 min at 4°C, resuspended in 200 μL of collagenase I solution (Sigma, C9722, 1 mg/mL in PBS), and digested at 25°C for 1 h with gentle trituration performed at 10 min intervals. The digestion was quenched by adding 1 mL of PBS containing 0.04% BSA (Thermo Fisher Scientific, AM2616). The cell suspension was washed, passed through a 40-μm Flowmi cell strainer (Bel-Art, H13680-0040), and centrifuged again at 500 × g for 5 min at 4°C. The pellet was resuspended in 50 μL of PBS with 0.04% BSA and subjected to gentle, repeated pipetting (∼200 cycles) to ensure complete dissociation into a single-cell suspension. Finally, the cell concentration and viability were assessed using a hemocytometer (Neubauer Improved, Optik Labor) under a Leica DM100 LED microscope.

### 10× genomics single-cell RNA sequencing and data processing

Single-cell RNA sequencing libraries were prepared from approximately 10,000 cells per sample using the Chromium Single Cell 3′ Library and Gel Bead Kit v3 (10× Genomics) according to the manufacturer’s protocol. The resulting libraries were sequenced on an Illumina NovaSeq 6000 platform. Raw sequencing data were processed with Cell Ranger software (v2.2.0) using the count and aggr workflows, aligned to the *Drosophila melanogaster* reference genome (FlyBase release r6.22). Cell calling was performed by the software based on barcode and UMI distributions. Two aggregated datasets were generated for downstream analysis: one from the wild-type dataset (*Cp*OGA^DM^), comprising 18,371 cells with a median of 1,284 genes per cell, and the other from the hypo-*O*-GlcNAcylation dataset (*Cp*OGA^WT^), comprising 15,613 cells with a median of 1,435 genes per cell. The aggregation was performed without normalization to preserve raw gene expression counts for subsequent analysis.

### Seurat data processing

The integrated dataset combining samples from the *Cp*OGA^DM^ (DM) and *Cp*OGA^WT^ (WT) conditions was processed using Seurat (v3.0). Initial quality control filtered out cells outside the range of 200-4,500 detected genes or with mitochondrial transcript percentages exceeding 20%, resulting in a final set of 18,371 DM cells and 15,613 WT cells for analysis. Gene expression counts were log-normalized using a scale factor of 10,000, and the 2,000 most highly variable genes were identified for downstream dimensionality reduction. Principal component analysis (PCA) was performed, and 16 principal components were selected for subsequent clustering based on the ElbowPlot, JackStraw, and PCHeatmap utilities. Cells were clustered using a graph-based algorithm with a resolution parameter of 0.8 and visualized in two dimensions using Uniform Manifold Approximation and Projection (UMAP).

### Sample preparation and electron microscopy

Wild-type *Drosophila melanogaster* third-instar larval brains were dissected and fixed overnight at 4°C in 2.5% glutaraldehyde (Macklin, G810415). After rinsing with fresh buffer, the samples were post-fixed in 2% osmium tetroxide (Electron Microscopy Sciences, 19150) for 1.5 h at room temperature. The tissues were then stained with 1% uranyl acetate (TED PELLA, 19481) overnight at 4°C, washed in double-distilled water, and dehydrated through a graded ethanol and acetone series (30–100%, Guoyao, 10000418). Subsequently, the samples were infiltrated and embedded in Epon 812 resin (Zhongjing Keyi, GS02660) and polymerized at 60°C for 48 h. Ultrathin sections (approximately 70 nm) were cut using a Leica ultramicrotome, collected on Pioloform-coated copper grids, and counterstained with lead citrate. Imaging was performed using a Hitachi transmission electron microscope operated at 80 kV. For quantitative analysis, the distance between the inner and outer nuclear membranes, the diameter of nuclear pore complexes (NPCs), and the mean electron density of the NPC central channel were measured using ImageJ software.

### Quantification and statistical analysis

All experiments were performed with at least three independent biological replicates. Statistical analyses were conducted using GraphPad Prism 8.0, applying specific tests as appropriate for each experimental design: two-tailed unpaired or paired t-tests for comparisons between two groups, and one-way ANOVA followed by Dunnett’s test for multiple comparisons. Data are presented as the mean ± SD, with significance levels denoted as follows: \**p* < 0.05, \*\**p* < 0.01, \*\*\**p* < 0.001; ns, not significant. KEGG pathway enrichment analysis of the *O*-GlcNAcome in neuroblast lineages was performed using the DAVID bioinformatics resource, and a protein-protein interaction network was constructed and analyzed using the STRING database.

## Acknowledgments

We gratefully acknowledge Drs. Ting Xie, Wu-Min Deng, Yikang Rong, Yang Yu, Guangshuo Ou, Xi Huang, Hai Huang, Zheng Guo, Yan Song, Sijun Zhu, Jiwu Wang, Bing Yang, Hansong Ma, Daan van Aalten, the Bloomington *Drosophila* Stock Center, and TsingHua Fly Center for inspiring discussions or reagents. This project has been supported by the National Natural Science Foundation of China (grants 32450591 and 32170821 to K.Y, and 32101034 to F.C), and Department of Science & Technology of Hunan Province (grants 2023RC1028 and 2023SK2091 to K.Y, and 2022JJ40762 to F.C).

## Author contributions

Conceptualization: F.C., D.v.A., X.Y., K.Y.;

Methodology: F.C., H.Y., W.Z., X.W., S.M., L.L., H.Q., K.L., H.H., K.Y.;

Validation: H.Y., W.Z., L.L.;

Software: F.C., S.M.;

Formal Analysis: F.C., H.Y., W.Z., S.M., K.Y.;

Investigation: F.C., H.Y., W.Z., X.W., S.M., L.L.;

Resources: K.L., H.H., D.v.A., Z.Z., X.Y., K.Y.;

DataCuration: F.C., H.Y., W.Z., S.M., K.Y.;

Writing-Original Draft: F.C., H.Y.;

Writing-Review & Editing: K.Y.;

Visualization: F.C., H.Y., W.Z., X.W., S.M., K.Y.;

Supervision: K.Y.;

Project Administration: L.L., K.Y.;

Funding Acquisition: K.Y.

## Supplementary Figures and Legends

**Extended Data Figure 1.**
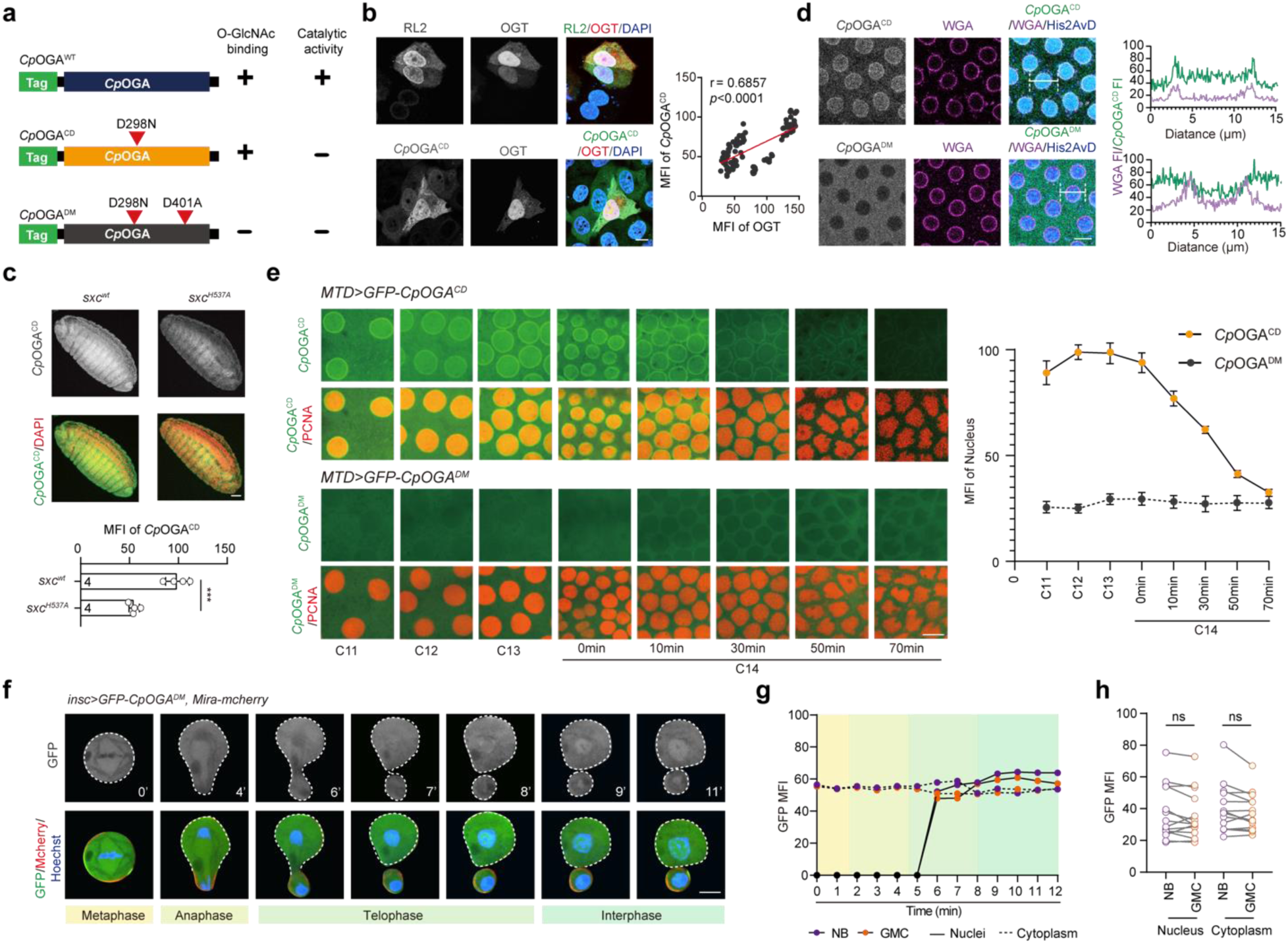
A fluorescently labeled mutant bacterial *O-*GlcNAcase for visualizing protein *O-*GlcNAcylation in living cells. **a.** Schematic of different versions of OGA from *Clostridium perfringens* (*Cp*OGA) fused to fluorescent tag. **b.** Representative images showing the Hela cells transfected with GFP-tagged human OGT (red). Anti-*O-*GlcNAc antibody RL2 (top row) or fluorescently labeled *Cp*OGA^CD^ protein (bottom row) is used to visualize the *O-*GlcNAcylated proteome (green). Nuclei are visualized with DAPI (blue). Correlation of mean fluoresecent intensity (MFI) of *Cp*OGA^CD^ staining versus GFP-OGT (right, n = 113). The Pearson correlation coefficient (r) and the *p-value* are shown. Scale bar, 10 μm. **c.** Representative images showing the *sxc* wild-type (*sxc^wt^*) and hypomorphic mutant (*sxc^H^*^537^*^A^*) embryos stained with GFP-*Cp*OGA^CD^ protein (green). DAPI staining is shown in red. Scale bar, 50 μm. Quantification of the MFI of *Cp*OGA^CD^ staining in *sxc^wt^* and *sxc^H^*^537^*^A^* embryos (bottom, n = 4). *P* value is determined by two-tailed unpaired t test, the stars indicate significant differences (\*\*\**p* < 0.001), error bars represent SD. **d.** Live imaging of GFP-*Cp*OGA^CD^ and GFP-*Cp*OGA^DM^ (green) injected into cycle-12 embryos. WGA is co-injected and shown in magenta, and His2AvD-RFP expressed from a transgene in blue. WGA (magenta) and the indicated *Cp*OGA fluorescent intensity (green) along the white lines are plotted on the right. Scale bar, 10 μm. **e.** Real-time imaging of early embryos expressing GFP-*Cp*OGA^CD^ and GFP-*Cp*OGA^DM^ (green). PCNA-mCherry protein is included in the injectant to visualize interphase nuclei. Embryonic cell cycles (C11, C12, C13) or relative times from the beginning of cycle 14 (in minutes) are shown on the bottom. Scale bar, 10 μm. Quantification of GFP-*Cp*OGA^CD^ and GFP-*Cp*OGA^DM^ MFI in the nuclei (right, n = 8). **f.** Real-time imaging of asymmetric cell division of NBs expressing GFP-*Cp*OGA^DM^ with Mira-Mcherry. Nuclei are stained with Hoechst (blue) and Mira are shown in red to label GMC. NBs and GMCs are outlined by white dashed lines. Times are indicated in minutes. Scale bar, 5 μm. **g.** Quantification of GFP MFI in the nuclei and cytoplasm of apical large cells (NBs) and basal small cells (ganglion mother cells, GMCs) over time. **h.** Quantification of nuclear and cytoplasmic GFP MFI during telophase of asymmetric cell division (n = 13). *P* values are determined by two-tailed paired t test. For the statistical data, the stars indicate significant differences (ns, not significant), error bars represent SD.

**Extended Data Figure 2.**
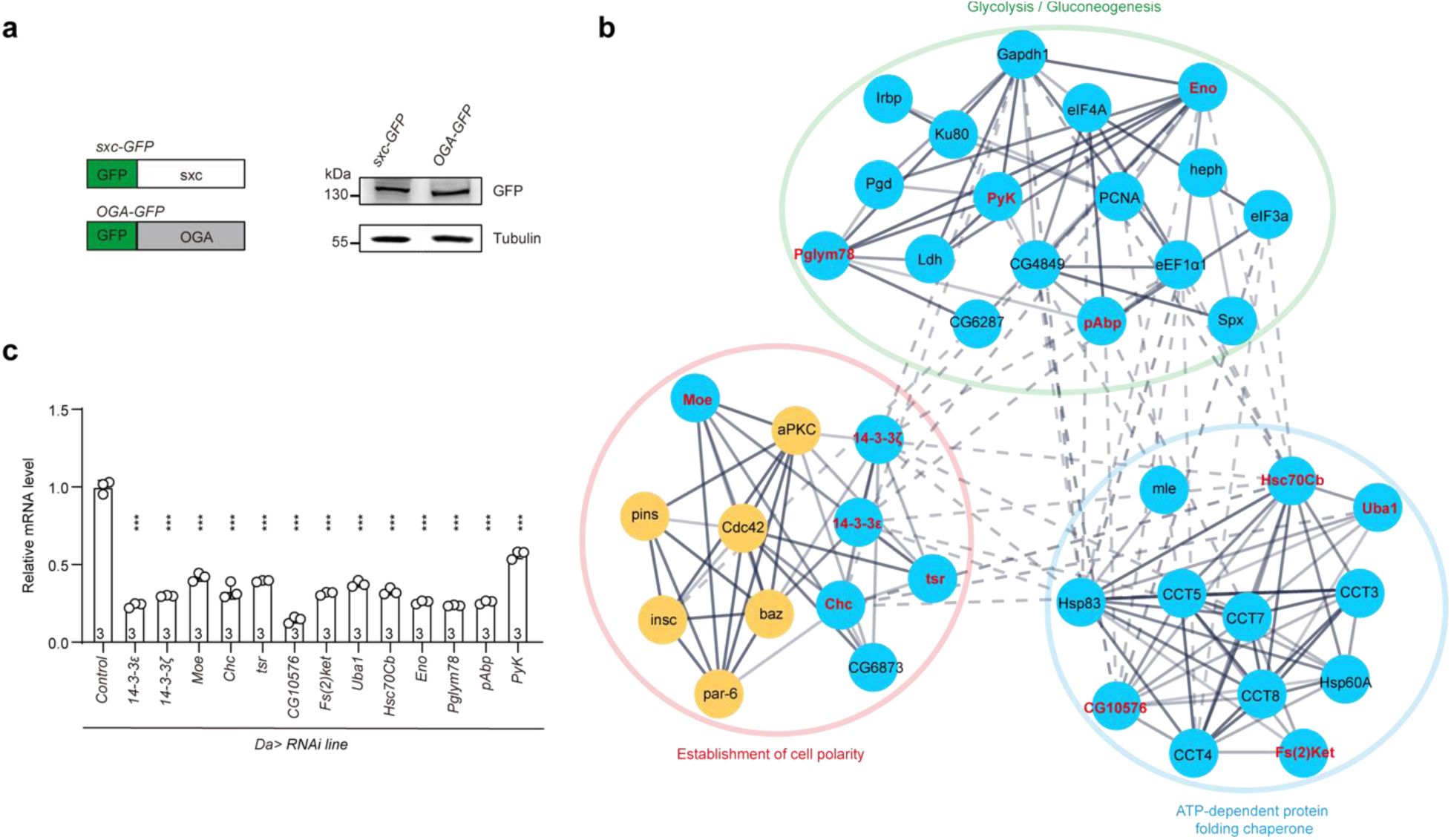
The screen of candidates that regulate the asymmetric distribution of the *O-*GlcNAcome. **a.** Schematic of sxc and OGA fused to GFP, and detection GFP-tagged sxc and OGA by western blot with anti-GFP antibody (right). **b.** STRING visualization of protein-protein interaction network, and the groups are color coded. **c.** qPCR analysis of the expression after shRNA-mediated knockdown. *P* values are determined by one-way ANOVA followed by Dunnett’s multiple comparison test and the stars indicate significant differences (\*\*\**p* < 0.001), error bars represent SD.

**Extended Data Figure 3.**
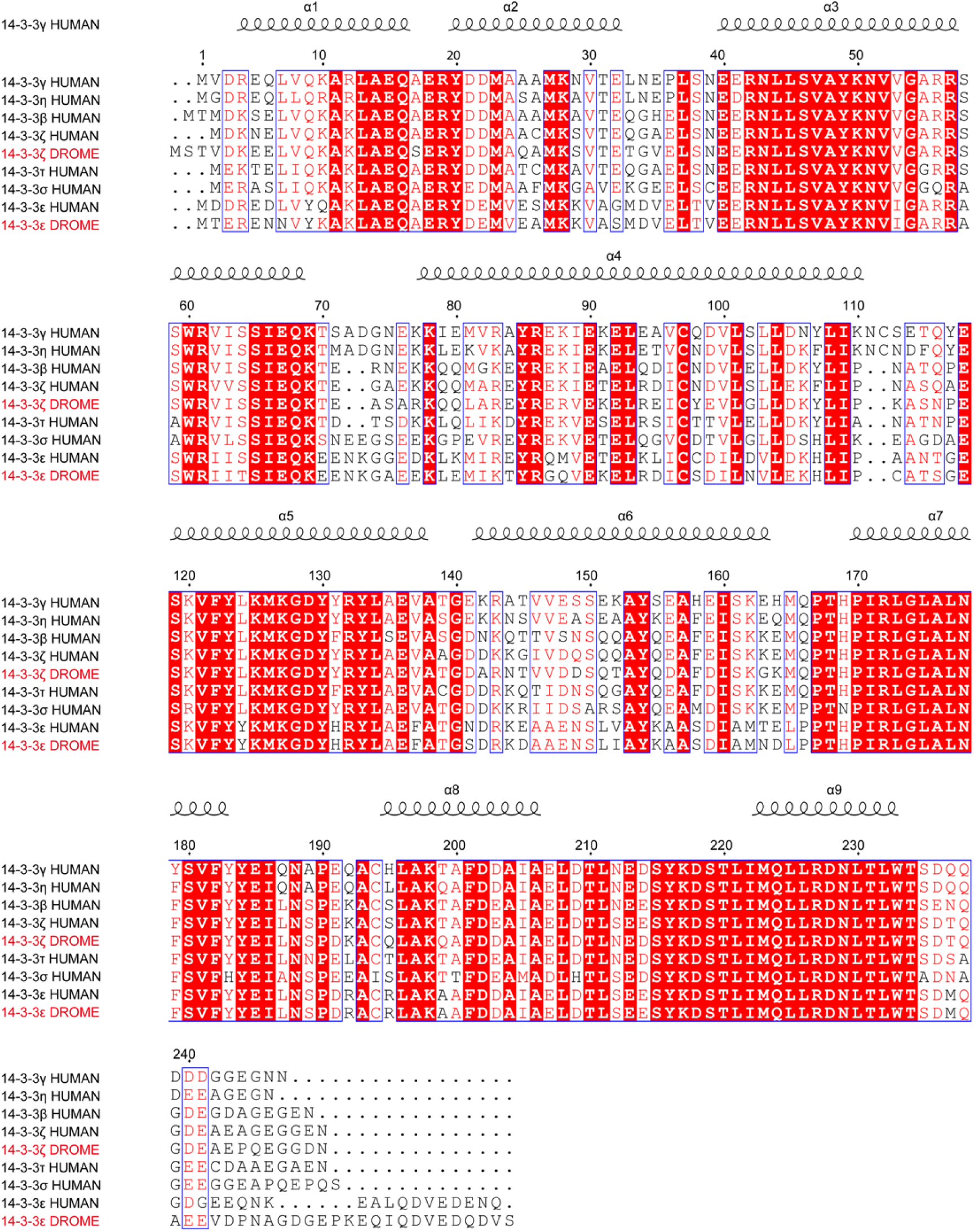
14-3-3 isoforms are conserved between humans and *Drosophila*. Protein sequence alignment of 14-3-3 isoforms from humans (7 isoforms) and *Drosophila* (2 isoforms). Residues identical in all sequences are printed in white on a red background, and residues identical or with a conservative substitution in at least seven of the nine sequences are printed in red on a white background.

**Extended Data Figure 4.**
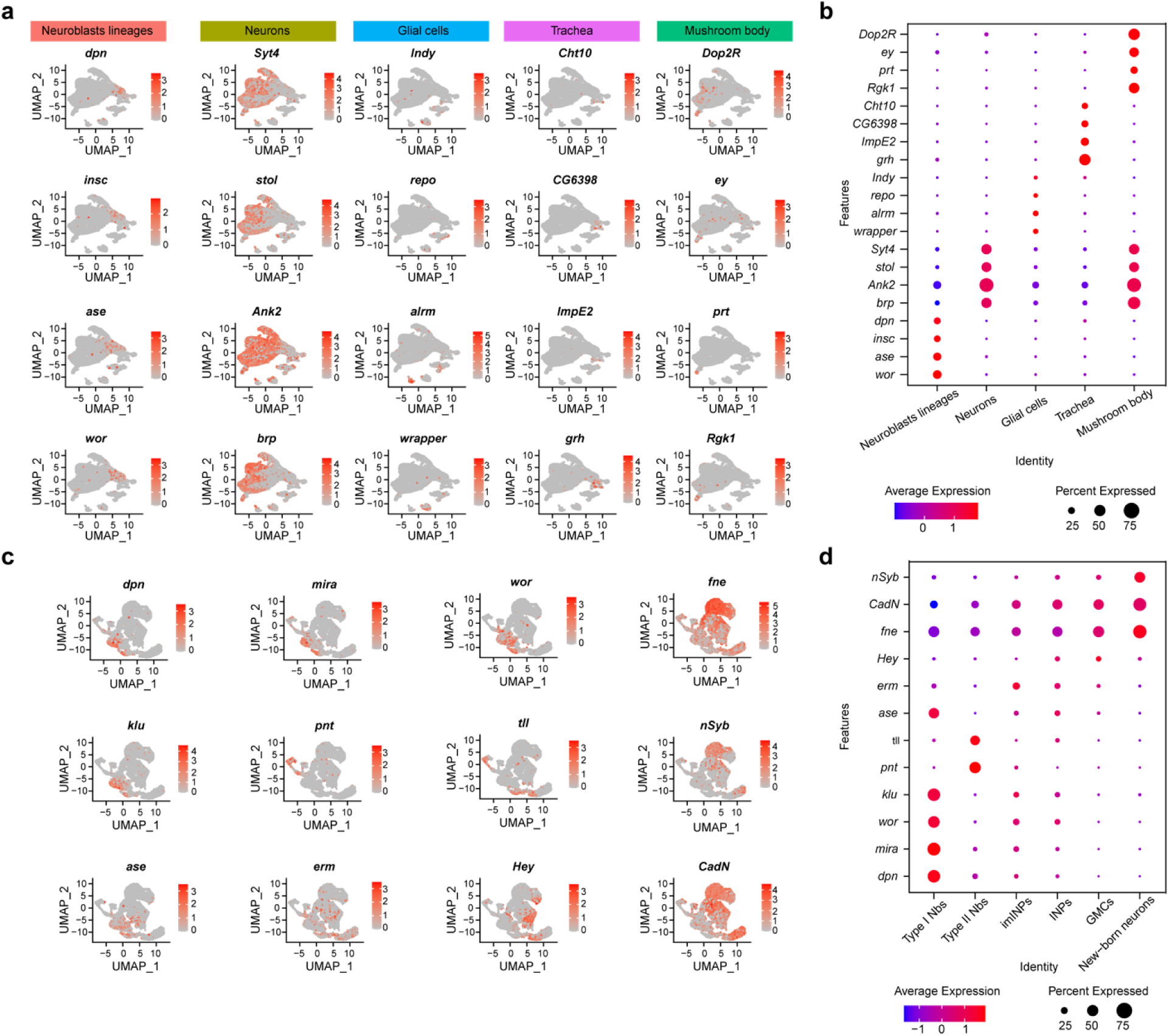
Characterization of cell composition of the larval brain and NB clones. **a, c.** Two dimensions UMAP plot labeling 5 subclusters marker genes of the larval brain (a) and NB clone marker genes (c). **b, d.** Bubble plot of the expression levels of the marker genes in the 5 subclusters of larval brain (b) and in the NB clones (d). Bubble size corresponds to the percentage of cells expressing a particular gene and color to gene expression intensity levels. Red: high expression, blue: low expression.

**Extended Data Figure 5.**
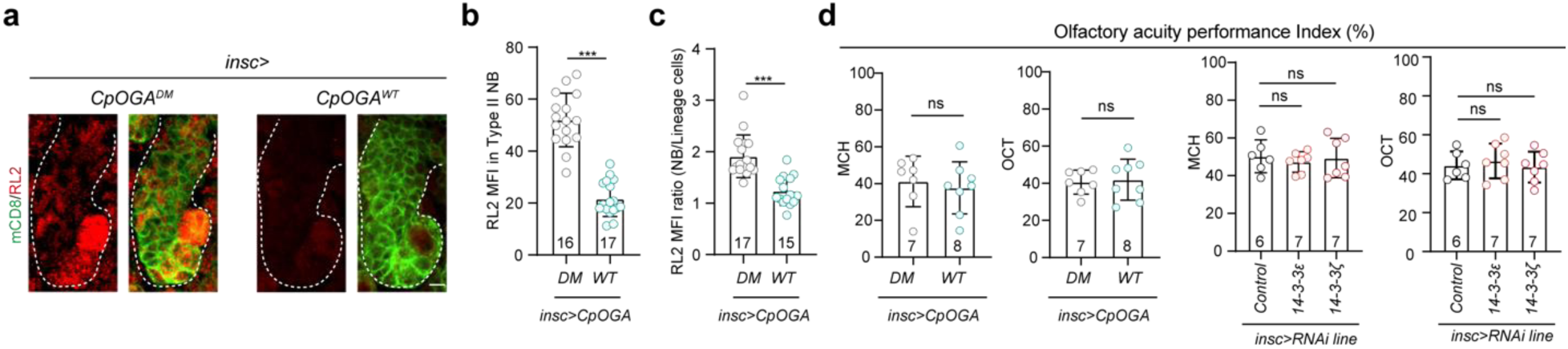
Interruption of the *O-*GlcNAcome homeostasis in NB lineage. **a.** Representative images of Type II NB clones expressing *Cp*OGA^WT^ or *Cp*OGA^DM^. *O*-GlcNAcome is visualized with RL2 (red) and the NBs and their lineage cells are marked with mCD8-GFP (green). The Type II NB clones are outlined by white dashed lines. Scale bar, 10 μm. **b.** Quantification of the RL2 MFI in NB clones (n = 16 and 17). **c.** Quantification of the ratio of RL2 MFI in NBs versus their lineage cells (n = 17 and 15). **d.** Bar graphs showing the odor acuity performance index of flies expressing *Cp*OGA^WT^ or *Cp*OGA^DM^, or with 14-3-3ζ or 14-3-3ε knockdown (n = 6 - 8). For the statistical data, *P* values in (b), (c), and first two bar graghs in (d) are determined by two-tailed unpaired t test. *P* values in the last two bar graphs in (d) are determined by one-way ANOVA followed by Dunnett’s multiple comparison test. The stars indicate significant differences (ns, not significant; \*\*\**p* < 0.001), error bars represent SD.

**Extended Data Figure 6.**
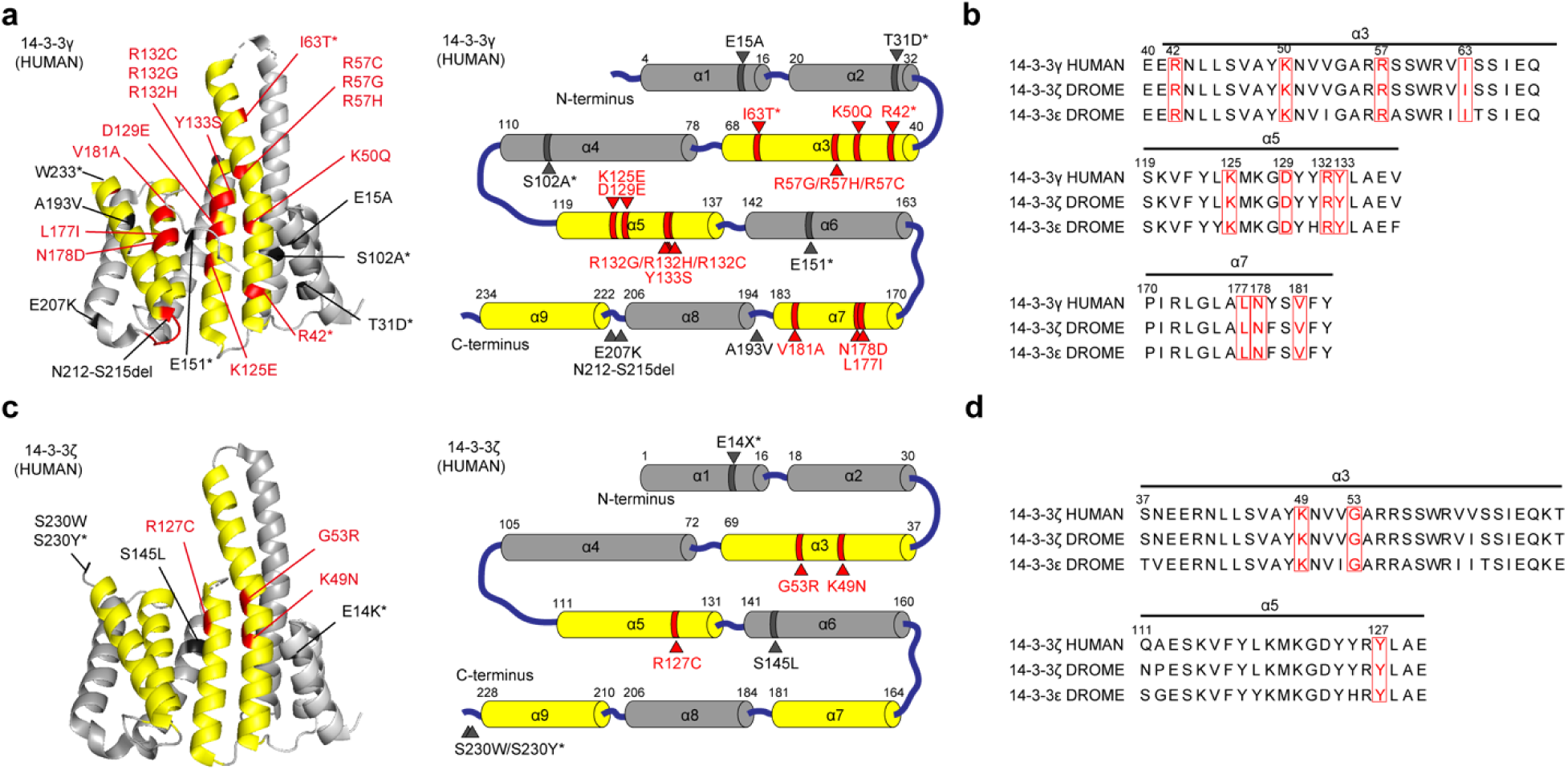
The pathogenic mutations in human 14-3-3γ and 14-3-3ζ are conserved in *Drosophila* isoforms. **a, c.** 3D crystal structure and pattern diagram with pathogenic mutational information of human 14-3-3γ (a) and 14-3-3ζ (c). The 14-3-3 protein consists of a bundle of conserved nine α-helices. The third, fifth, seventh, and ninth helices (yellow) form an amphipathic groove that interacts with target proteins. Pathogenic mutations within amphipathic groove are marked in red, while others are marked in black. **b, d.** Pathogenic mutations (red) located within the amphipathic groove of human 14-3-3ζ are conserved in *Drosophila* 14-3-3ζ and 14-3-3ε.

**Extended Data Figure 7.**
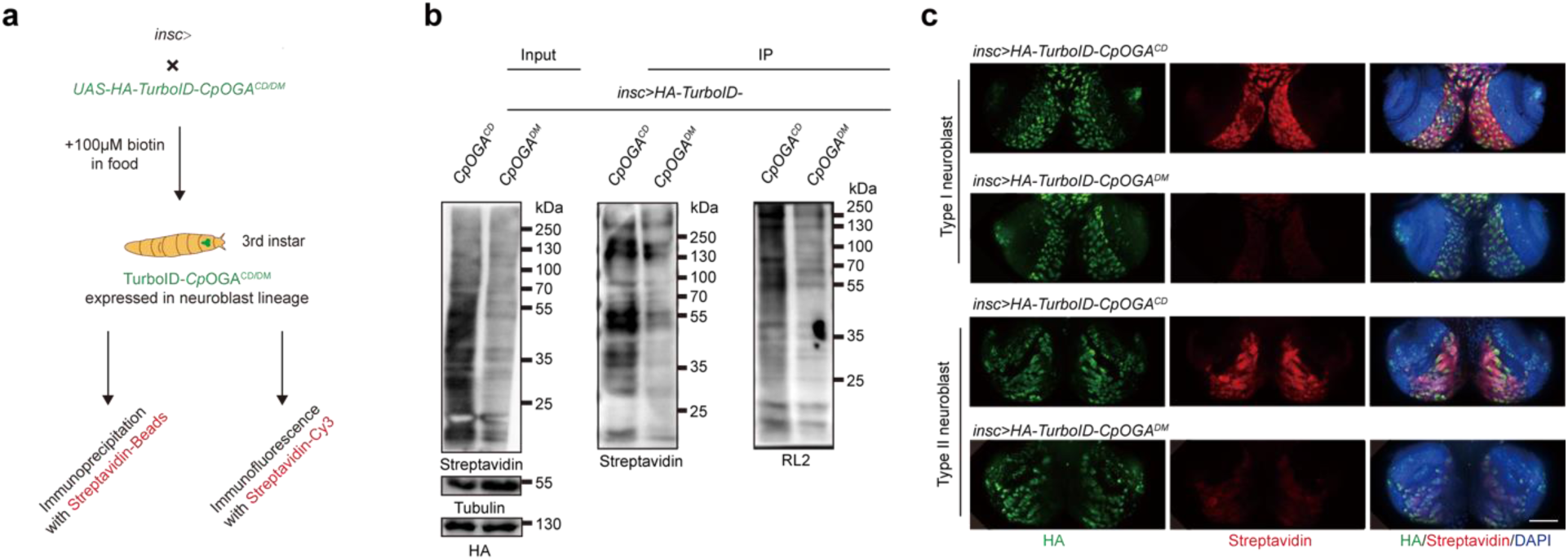
Analysis of tissue-specific *O-*GlcNAcylated proteins in NBs and their lineages using TurboID-*Cp*OGA^CD^. **a.** Scheme for the validation of the TurboID-*Cp*OGA^CD^ system in larvae brain. **b.** Immunoprecipitation of biotinylated proteins from larvae brain. Biotinylation is detected by immunoblotting with streptavidin-HRP, and *O-*GlcNAcylation with anti-*O-*GlcNAc antibody (RL2). The expression of TurboID-*Cp*OGA^CD/DM^ is validated by anti-HA immunoblotting. **c.** Representative images of the larval brain lobes. Biotinylated proteins are stained with streptavidin-Cy3 (red), and TurboID-*Cp*OGA^CD^ with anti-HA antibody. Nuclei are visualized by DAPI (blue). Scale bar, 100 μm.

**Extended Data Figure 8.**
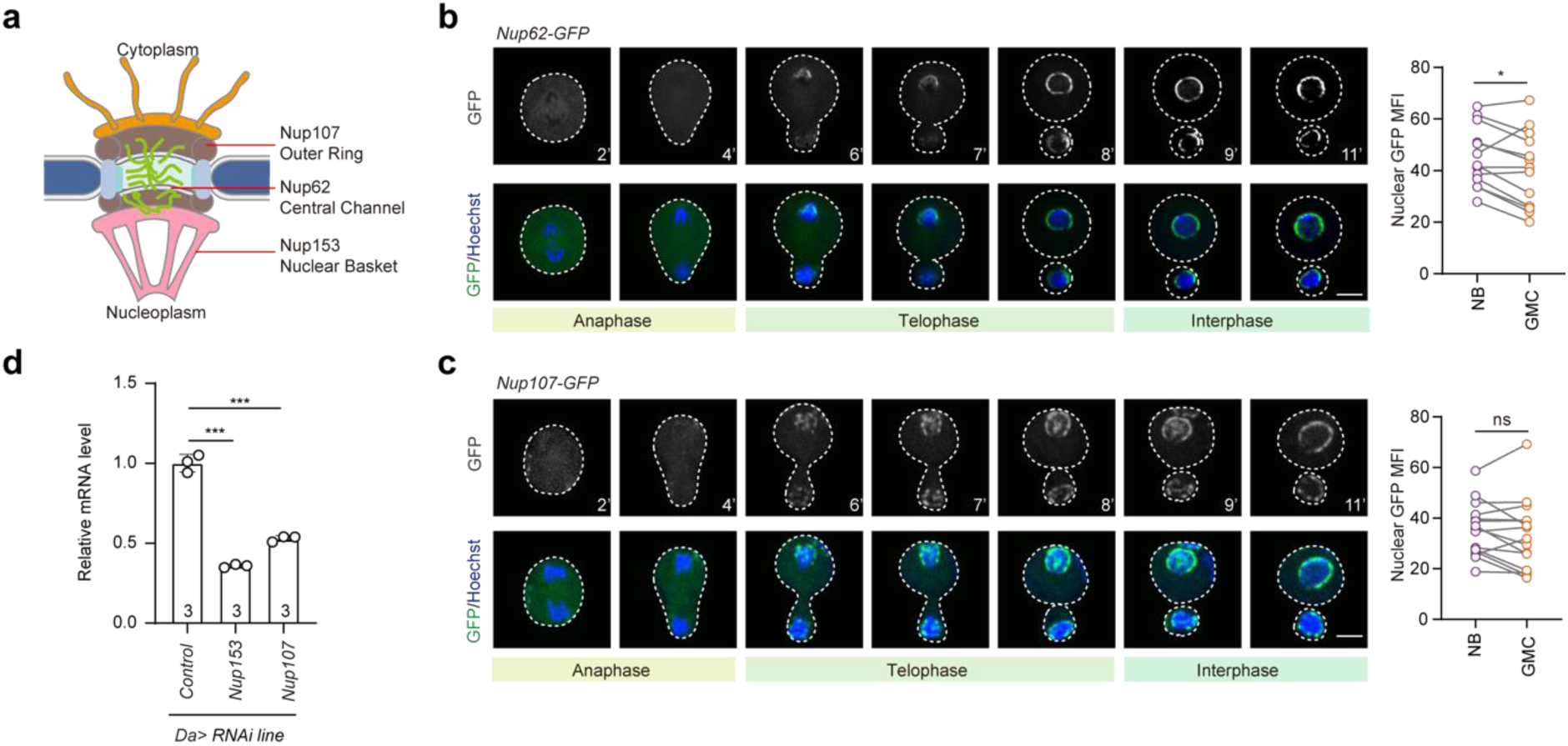
Symmetric distribution of nuclear pore proteins Nup107 and NUP62 during the asymmetric division of NBs. **a.** Scheme of nuclear pore structure, in which representative *O-*GlcNAcylated nuclear pore proteins of different components are labeled. **b-c.** Real-time imaging of asymmetric cell division of NBs with Nup62-GFP knockin (c) or Nup107-GFP knockin (d). Nuclei are stained with Hoechst (blue). Elapsed times are indicated in minutes. NBs and GMCs are outlined by white dashed lines. Scale bars, 5 μm. Quantification of nuclear GFP MFI during telophase of asymmetric cell division (n = 13 - 15). *P* values are determined by two-tailed paired t test. For the statistical data, the stars indicate significant differences (ns, not significant; \**p* < 0.05). **d.** qPCR analysis of knockdown efficiencies of *Nup153* and *Nup107*. *P* values are determined by one-way ANOVA followed by Dunnett’s multiple comparison test. The stars indicate significant differences (\*\*\**p* < 0.001), error bars represent SD.

**Extended Data Figure 9.**
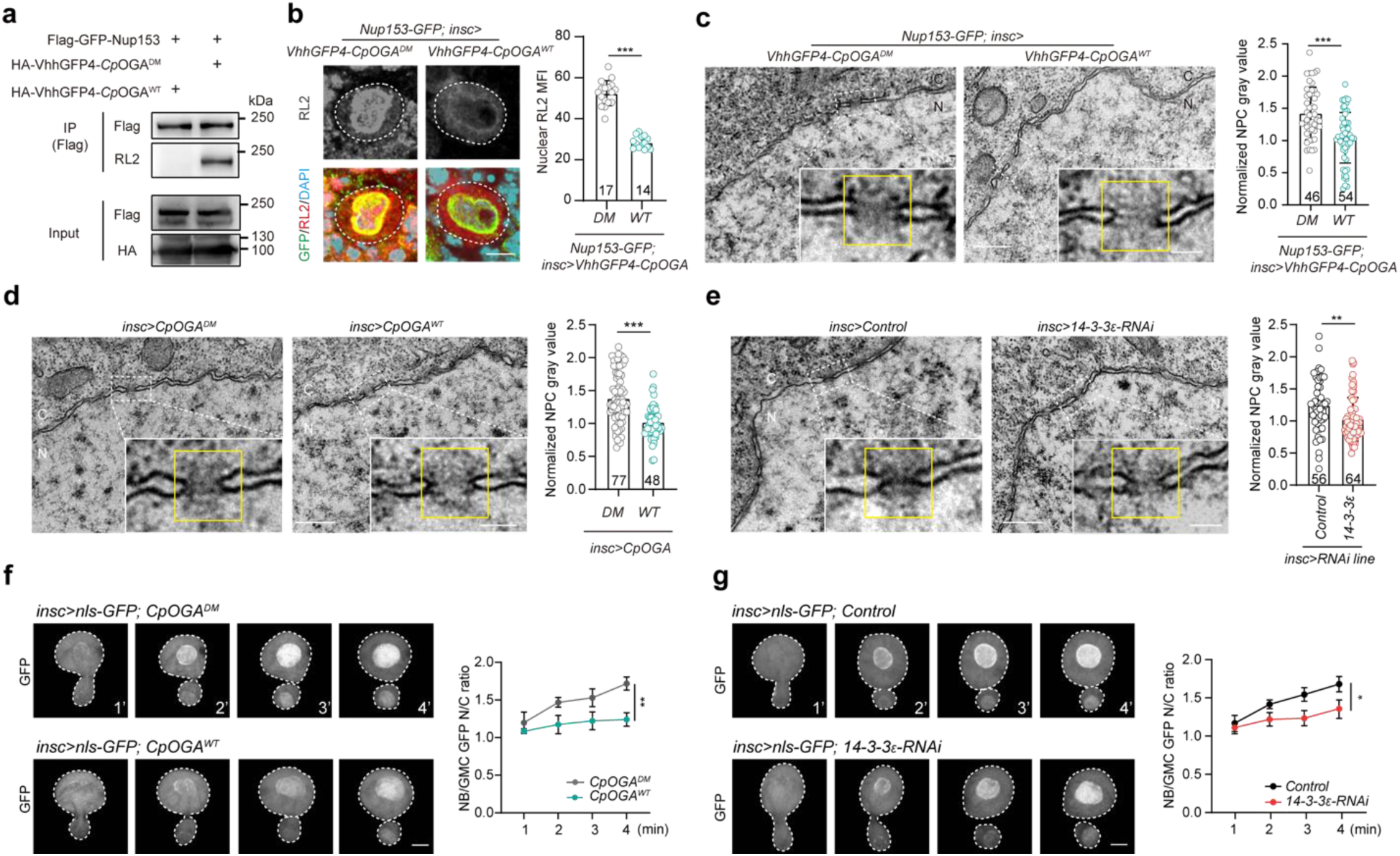
*O-*GlcNAcylation regulates nuclear pores and nucleocytoplasmic transport in neuroblasts. **a.** Validation of the targeted de-*O*-GlcNAcylation strategy using the Flag-GFP-tagged Nup153. The Flag-GFP-Nup153 is co-expressed with HA-tagged GFP nanobody-*Cp*OGA^WT^ (HA-VhhGFP4-*Cp*OGA^WT^) or its control HA-VhhGFP4-CpOGA^DM^. The immunoprecipitated Flag-GFP-Nup153 is blotted with RL2 antibody to detect *O*-GlcNAcylation. **b.** Representative images of Nup153-GFP knockin neuroblasts expressing GFP nanobody-*Cp*OGA^WT^ (VhhGFP4-*Cp*OGA^WT^) or its control VhhGFP4-*Cp*OGA^DM^. *O*-GlcNAcylation level is assessed with RL2 staining (red), and nuclei are stained with DAPI (blue). Scale bar, 5 μm. Quantification of nuclear RL2 MFI is shown on the right (n = 14 and 17). **c.** Representative TEM images showing mature NPCs (nuclear pore complex, indicated by yellow box) in NBs of the indicated genotypes. N, nucleus; C, cytoplasm. Scale bars, 500 nm (left), 100 nm (right). Quantification of the normalized electronic density value within the NPCs (right, n = 46 and 54). **d-e.** Representative TEM images showing mature NPCs in NBs of indicated genotypes. N, nucleus; C, cytoplasm. Scale bars, 500 nm (left), 100 nm (right). Quantification of normalized NPC gray value (right, n = 48 - 77). **f-g.** Real-time imaging analysis of asymmetric division of NBs expressing nls-GFP with *Cp*OGA^WT^/*Cp*OGA^DM^ overexpression or 14-3-3ε knockdown driven by *insc-Gal4*. Elapsed times are indicated in minutes. NBs and GMCs are outlined by white dashed lines. Scale bars, 5 μm. Quantification of relative nuclear GFP MFI from telophase of asymmetric cell division to interphase (right, n = 4). For statistical analyses, *P* values in (b), (c), (d), (e) are determined by two-tailed unpaired t test, while those in (f), (g) are determined by two-way ANOVA. The stars indicate significant differences (\**p* < 0.05, \*\**p* < 0.01, \*\*\**p* < 0.001), error bars represent SD.

## Supplementary Videos

**Supplementary Video 1**

Real-time imaging of asymmetric cell division of NBs expressing GFP-*Cp*OGA^CD^ (left) or GFP-*Cp*OGA^DM^ (right). NBs and GMCs are outlined by white dashed lines. Times are indicated in minutes. Scale bars, 5 μm.

**Supplementary Video 2**

Real-time imaging of symmetric cell division of GMCs expressing GFP-*Cp*OGA^CD^. Progeny cells are outlined by white dashed lines. Nuclei are stained with Hochest (blue). Times are indicated in minutes. Scale bars, 5 μm.

## Supplementary Tables

**Supplementary Table 1**

List of mammalian *O*-GlcNAc reader proteins and their homologs in *Drosophila*

**Supplementary Table 2**

TM score of topological similarity comparison between human 14-3-3 and *Drosophila* 14-3-3

**Supplementary Table 3**

14-3-3 related disease mutation site information

**Supplementary Table 4**

*O*-GlcNAcylated proteins identified by TurboID-CpOGA^CD^ from neuroblast lineage of *Drosophila*

**Supplementary Table 5**

KEGG analysis of *O*-GlcNAc proteins in the *Drosophila* neuroblast lineage

**Supplementary Table 6**

Currently reported *O*-GlcNAc information of nucleoporins

**Supplementary Table 7**

Sequences of all the primers used in this study

